# Regulation of the Dimerization and Activity of SARS-CoV-2 Main Protease through Reversible Glutathionylation of Cysteine 300

**DOI:** 10.1101/2021.04.09.439169

**Authors:** David A. Davis, Haydar Bulut, Prabha Shrestha, Amulya Yaparla, Hannah K. Jaeger, Shin-ichiro Hattori, Paul T. Wingfield, Hiroaki Mitsuya, Robert Yarchoan

## Abstract

SARS-CoV-2 encodes main protease (M^pro^), an attractive target for therapeutic interventions. We show M^pro^ is susceptible to glutathionylation leading to inhibition of dimerization and activity. Activity of glutathionylated M^pro^ could be restored with reducing agents or glutaredoxin. Analytical studies demonstrated that glutathionylated M^pro^ primarily exists as a monomer and that a single modification with glutathione is sufficient to block dimerization and loss of activity. Proteolytic digestions of M^pro^ revealed Cys300 as a primary target of glutathionylation, and experiments using a C300S M^pro^ mutant revealed that Cys300 is required for inhibition of activity upon M^pro^ glutathionylation. These findings indicate that M^pro^ dimerization and activity can be regulated through reversible glutathionylation of Cys300 and provides a novel target for the development of agents to block Mpro dimerization and activity. This feature of M^pro^ may have relevance to human disease and the pathophysiology of SARS-CoV-2 in bats, which develop oxidative stress during flight.

## INTRODUCTION

The main protease (M^pro^) of SARS-CoV-2 coronavirus is encoded as part of two large polyproteins, pp1a and pp1ab, and is responsible for at least 11 different cleavages. Thus, M^pro^ is essential for viral replication and has been identified as a promising target for the development of therapeutics for treatment of coronavirus disease 2019 (COVID-19) ^1,2^. M^pro^ is known as a 3C-like protease (3CL) due to its similarity to picornavirus 3C protease in its cleavage site specificity ^3^. Through extensive studies on M^pro^ from SARS-CoV-1, whose sequence is 96% identical to SARS-CoV-2 M^pro^, a wealth of information has been obtained that can be applied to studies now ongoing with SARS-CoV-2 M^pro^ (for review see ^4^). M^pro^ of SARS-CoV-1 and SARS-CoV-2 consist of three major domains, I, II, and III. Unlike other 3C-like proteases, studies on M^pro^ from SARS-CoV-1 and SARS-CoV-2 have revealed that they are only active as homodimers even though each individual monomeric subunit contains its own active site ^5,6^. Studies on SARS-CoV-1 to explain why dimerization is required for activity have revealed that, in the monomeric state, the active site pocket collapses and is not available for substrate binding and processing ^7^. In these studies it was also revealed that the extra domain (III) plays a key role in dimerization and activation of M^pro^ and that arginine 298 in this domain is essential to allow proper dimerization and M^pro^ activity ^7^.

The proteases of HIV and other retroviruses are also active as homodimers, and we previously demonstrated that each of the retroviral proteases studied (HIV-1, HIV-2 and HTLV-1) could be reversibly regulated through oxidation of residues involved in protease dimerization ^8,9,10,11^. The activity of HIV-1 and HIV-2 protease can be reversibly inhibited by oxidation of residue 95, located at the dimer interface ^9^ and these oxidative modifications are reversible with cellular enzymes, glutaredoxin (Grx) and/or methionine sulfoxide reductase, respectively ^12,13^. The majority of other retroviral proteases also have one or more cysteine and/or methionine residues at the dimer interface region and modification of these residues, under conditions of oxidative stress, would be predicted to similarly regulate dimerization and activity ^8^. There is further evidence that HIV polyprotein precursors encoding these proteases are initially formed in an oxidized inactive state and need to be activated in a reducing environment ^8,9,13,14,15^ Moreover, the initial step in HIV-1 polyprotein processing, which is required to release the mature protease, is also regulated through reversible oxidation of cysteine 95 ^16^.

In addition to the active site cysteine, M^pro^ of SARS-CoV-1 and SARS-CoV-2 contain 11 other cysteine residues throughout the 306 amino acid sequence and all these residues are present in their reduced form in the crystal structures of M^pro^. This is a relatively large number of cysteines for a protein of this size (3.9% vs 2.3% average cysteine content of human proteins) ^17^. While a number of the cysteines are buried and may not be exceptionally susceptible to oxidation in the native structure, there are certain cysteine residues (notably cysteine 22, 85, 145, 156 and 300) that are at least partially surface/solvent exposed and potentially susceptible to oxidative modification. Here, we demonstrate that dimerization and activity of SARS-CoV-2 M^pro^ can be regulated through reversible glutathionylation of cysteine 300.

## RESULTS

### Treatment of M^pro^ with oxidized glutathione inhibits protease activity

M^pro^ activity was measured utilizing a para nitroanilide (pNA) substrate (H2N-TSAVLQ-pNA) as described previously for SARS-CoV-1 M^pro18^. To assess the effects of oxidized glutathione (GSSG) and reduced glutathione (GSH) on M^pro^, we treated M^pro^ at concentrations of either 1.2 or 18 µM with 2 mM or 10 mM of GSSG or GSH for 30 minutes at 37°C and then measured activity. Previous reports have indicated that the K_d_ of M^pro^ dimerization is about 2 µM ^6^ and that is consistent to what we found in this work. Thus, M^pro^ would be predicted to be largely monomeric at 1.2 µM and dimeric at 18 µM. After exposure of 1.2 µM M^pro^ to 2 mM GSSG, activity was inhibited by an average of 44% while after exposure to 10 mM GSSG, activity was inhibited by more than 90% (Figure 1A). By contrast, GSH had little effect or somewhat increased protease activity at these concentrations (Figure 1A). Interestingly, when the M^pro^ concentration was increased to 18 µM it was largely resistant to GSSG inhibition, with no inhibition observed with 2 mM GSSG and less than 20% inhibition with 10 mM GSSG (Figure 1B). These results suggest that monomeric M^pro^ may be more sensitive to glutathionylation than dimeric M^pro^. To confirm that M^pro^ was becoming modified with glutathione under these conditions, we acidified the samples at the end of the enzyme assays with formic acid/trifluoroacetic acid (FA/TFA) to arrest activity and glutathionylation and analyzed them by RP-HPLC/MALDI-TOF. The extent of glutathionylation was assessed by determining the mass of M^pro^ by protein deconvolution and by looking for the addition of approximately 305 amu and/or multiples of 305 to M^pro^ consistent with the addition of glutathione(s) via a disulfide bond. As revealed by RP-HPLC/MALDI-TOF analysis, treatment of 1.2 µM M^pro^ with 2 mM GSSG led to an estimated 45% monoglutathionylation (only an estimate based on the mass abundances), whereas treatment with 10 mM GSSG led to mono- (11%), di- (50%), and tri-glutathionylation (35%), with less than 4% of M^pro^ remaining unmodified (Figure 1C). Comparing the results of Figure 1A and 1C, the loss of M^pro^ activity correlated with the extent of glutathionylation. Interestingly, the data obtained with 2 mM GSSG suggested that modification of only one cysteine may be sufficient to lead to inhibition of M^pro^ activity, as this treatment yielded about 45% monoglutathionylation and showed an average 40% decrease in activity. By contrast, M^pro^ incubated at 18 µM during treatment with 2 mM GSSG showed very little modification or reduction in activity (Figures 1B and 1D). Moreover, treatment of 18 µM M^pro^ with 10 mM GSSG led to only 14% monoglutathionylation (Figure 1D), which was associated with an average inhibition of 18% (Figure1B).

**Figure 1:**
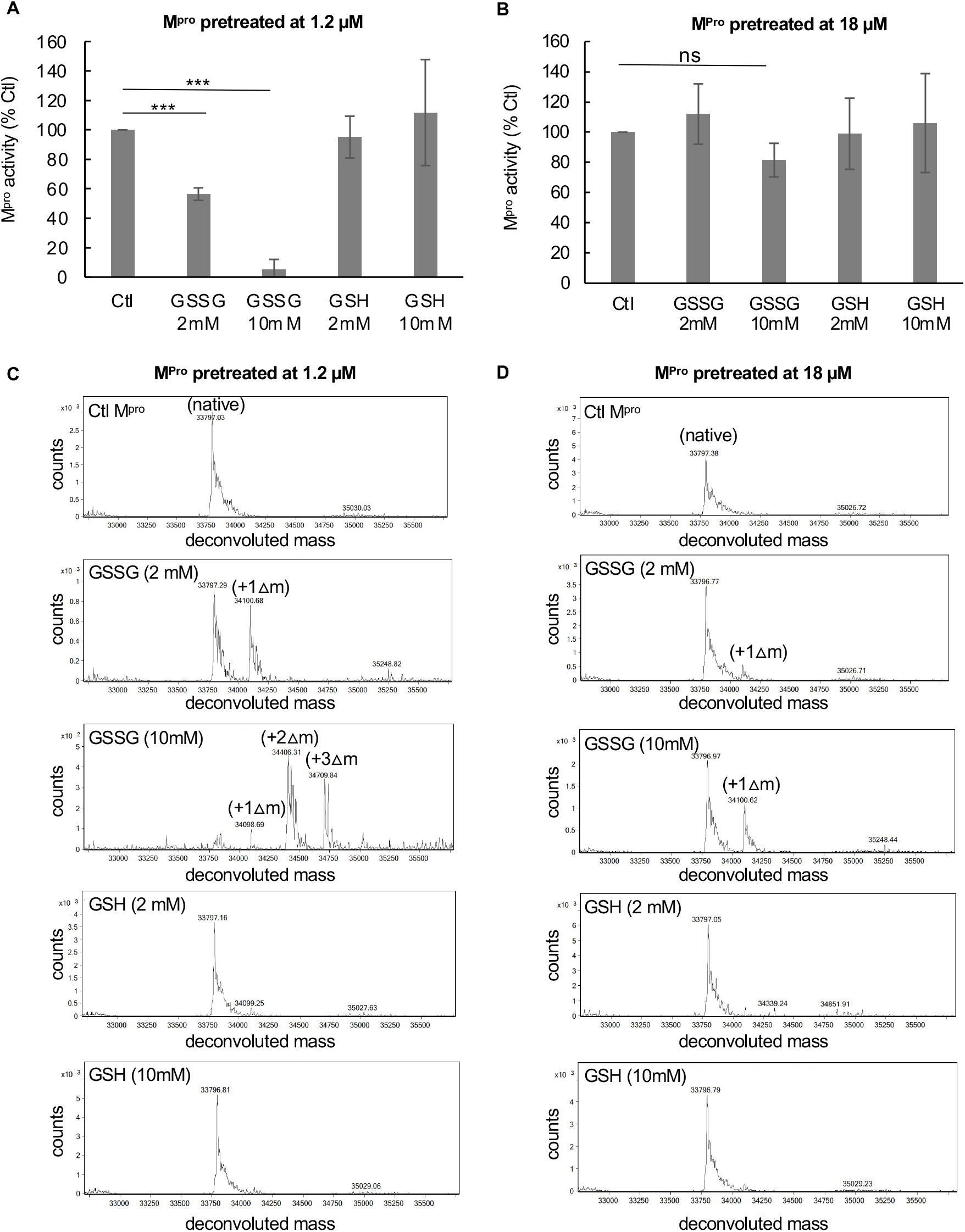
Exposure of low concentrations of SARS-CoV-2 M^pro^ to oxidized glutathione results in glutathionylation and inhibition of activity. (A,B) Activity of M^pro^ following a 30-minute pre-incubation of (A) 1.2 µM M^pro^ or (B) 18 µM M^pro^ pretreated with 2 mM or 10 mM oxidized or reduced glutathione. After preincubation, M^pro^ was assayed for protease activity at an equal final enzyme concentration (1 µM). (C,D) M^pro^ molecular masses found by protein deconvolution for M^pro^ eluting off of the C18 reverse phase column following the different treatments at (C) 1.2 µM and (D) 18 µM. The theoretical molecular mass of M^pro^ is 33796.48 and the deconvoluted molecular mass for controls in (C) and (D) was 33797.09 and 33,797.34, respectively, as determined using Agilent’s Mass Hunter software. The experimental masses are shown above each peak obtained by deconvolution. The native M^pro^ as well as the increases in masses indicative of glutathionylation are indicated for the addition of 1 (+Δ1), 2 (+Δ2), and 3 (+Δ3) glutathione moieties in the deconvolution profiles of GSSG-treated M^pro^. Observed increases were 304, 609, and 913 as compared to the predicted increases of 305.1, 610.2 and 915.3 for addition of 1, 2 or 3 glutathione’s, respectively. Based on the abundances, the estimated percent of monoglutathionylation in (C) at 2 mM GSSG was 45% and for 10 mM GSSG there was an estimated 11% mono, 50% di, and 35% tri-glutathionylation, respectively. In (D) after treatment with 2 mM GSSG there was <5% monoglutathionylation and after 10 mM GSSG there was an estimated 34% monoglutathionylation. For (A) and (B) the values shown are the mean and standard deviation for three independent experiments (n=3) while for (C) and (D) the analysis was one time. (*** = p-value < 0.005, paired Students *t-test*). All other comparisons to control activity were not found to be significant p-value >0.05). M^pro^ control activity for (A) was 6.42 +/− 2.5 µM/min/mg and for (B) was 9.6 µM/min/mg, and the percent activity in the treatments is normalized to their respective controls.

### Inhibition of M^pro^ activity by glutathionylation is reversible

To better understand the nature of M^pro^ inhibition by glutathionylation, we modified M^pro^ with 10 mM GSSG at pH 7.5, as described in the Materials and Methods, so that nearly all the M^pro^ was modified with at least one glutathione. Excess GSSG was removed by washing through an Amicon 10 kDa cut-off membrane. RP/HPLC//MALDI-TOF analysis of this preparation on a C18 column followed by protein deconvolution indicated M^pro^ was now a mixture of mono (23%), di (68%) and triglutathionylated forms (9%) with little detectable unmodified M^pro^ (based on abundances form protein deconvolution) (Figure 2A). To determine if the modification was reversible with thiol reducing agents, we treated the glutathionylated preparation with 10 mM DTT for 30 minutes. This resulted in more than 90% of the glutathionylated M^pro^ being converted back to native M^pro^ (Figure 2B). We then tested the activity of glutathionylated M^pro^. Glutathionylated M^pro^ had less than 5% of the activity of unmodified M^pro^, confirming that glutathionylation was inhibiting protease activity (Figure 2C). Following the addition of 10 mM DTT, the activity was fully restored, while DTT marginally improved native M^pro^ activity (Figure 2C).

**Figure 2:**
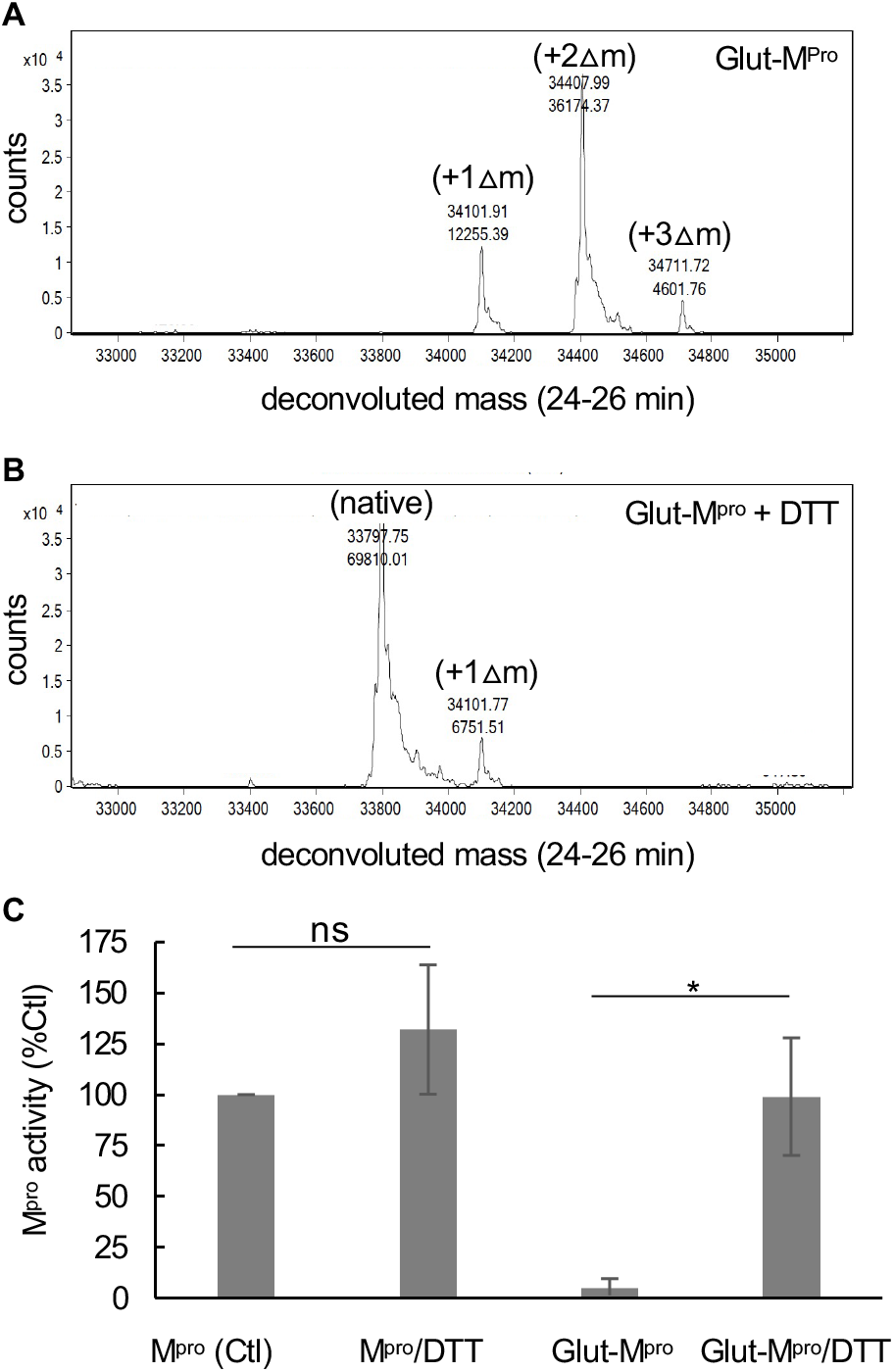
Inhibition of M^pro^ by glutathionylation can be reversed with reducing agent. M^pro^ was glutathionylated at pH 7.5 with 10 mM GSSG as described in the Materials and Methods and the extent of glutathionylation was determined by RP/HPLC/MALDI-TOF using a 2% acetonitrile gradient as described in materials and Methods. (A,B) Deconvoluted masses obtained by protein deconvolution of the M^pro^ peak (eluting between 24 and 26 min) for (A) 3 µg (5 µL injection) purified glutathionylated M^pro^ and (B)) 3 µg (5 µL injection) glutathionylated M^pro^ after a 30 min treatment with 10 mM DTT. Shown above each peak is the molecular mass (top number) and the abundance (bottom number) found by protein deconvolution. The native, monoglutathionylated (+Δ1), diglutathionylated (+Δ2), and triglutathionylated (+Δ3) M^pro^, are indicated in the figures. (C) M^pro^ activity (1 µM final enzyme) for native and glutathionylated M^pro^ preparations after a 30-min incubation in the absence or presence of 10 mM DTT. M^pro^ activity for control in (C) and was 4.95 +/− 1.2 µM/min/mg and percent activity for the different conditions was normalized to their respective controls. The values shown are the average and standard deviation from three separate experiments (n=3) (* = p-value < 0.05, paired students *t-test*, ns = not significant).

### Glutathionylation of M^pro^ inhibits M^pro^ dimerization

To assess M^pro^ dimerization we established a method consisting of size exclusion chromatography (SEC) coupled to mass spectrometry similar to that described previously for HIV-1 protease ^14^. In the SEC experiments we initially used SEC3000 columns and later SEC2000 columns from Phenomenex, both which could be used successfully to separate M^pro^. When injected at 60 µM on a SEC3000 column, unmodified M^pro^ eluted at 8.8 minutes (Figure 3A, black tracing) while glutathionylated M^pro^ eluted at 9.2 minutes (Figure 3A, red tracing). Protein deconvolution of the eluting M^pro^ confirmed the expected mass for unmodified M^pro^ (Figure 3B, black) and the glutathionylated forms of M^pro^ (Figure 3C, red). However, when injected at 7.5 µM, unmodified M^pro^ clearly eluted as two peaks at 8.9 and 9.4 minutes (Figure 3D, black tracing), while the glutathionylated M^pro^ still predominantly eluted at the later retention time (9.4 minutes) (Figure 3D, red tracing). Again, the masses for native and glutathionylated M^pro^ were confirmed (Figure 3E black tracing and 3F red tracing, respectively). Thus, the unmodified M^pro^ appeared to behave as a typical monomer/dimer two-species system with dimerization dependent on concentration, while glutathionylated M^pro^ behaved essentially as a single monomer-like species independent of its concentration. We carried out equilibrium analytical ultracentrifugation (AUC) on M^pro^ and glutathionylated M^pro^ to obtain both the molecular mass of the species and the K_d_ for dimerization. Matched native and glutathionylated M^pro^ samples (18 µM) were analyzed by AUC. The results indicated that native M^pro^ was in equilibrium between monomeric and dimeric forms and behaved with a calculated dimerization K_d_ of 2.4 µM (Figure 3G); consistent with previous reports ^6^. At high concentrations (60 µM), it was almost completely dimeric. By contrast, under the same conditions, the glutathionylated M^pro^ behaved almost completely monomeric with an estimated K_d_ of 200 µM (Figure 3H), indicating that glutathionylation was inhibiting dimerization of M^pro^.

**Figure 3:**
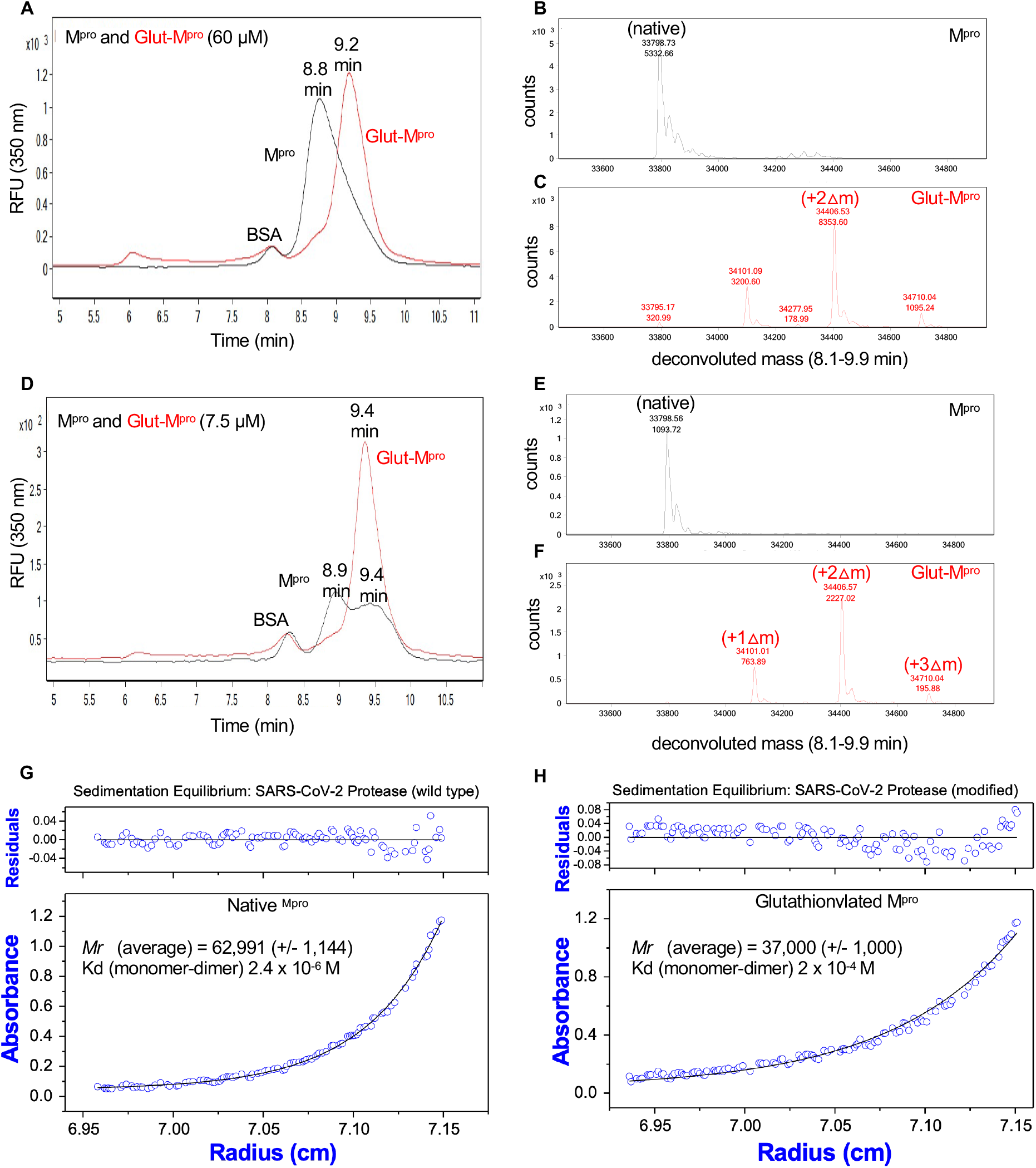
Size exclusion chromatography and equilibrium analytical ultracentrifugation of M^pro^ and glutathionylated M^pro^ indicates glutathionylated M^pro^ behaves as a monomer. (A,D) M^pro^ and glutathionylated M^pro^ were analyzed by SEC3000/MALDI-TOF. (A) Overlay of the chromatograms for 60 µM each of M^pro^ (black line) and glutathionylated M^pro^ (red line) and (D) 7.5 µM each of M^pro^ (black line) and glutathionylated M^pro^ (red line) by monitoring the intrinsic protein fluorescence (excitation 276 nm, emission 350 nm). Glutathionylated M^pro^ was made with 10 mM GSSG at pH 7.5 for 2-2.5 hours as described in Materials and Methods. (C,D) Protein deconvolution profiles for (B) native M^prov^ and (C) glutathionylated M^pro^ that were run as shown in (A). (E,F) Protein deconvolution profile for (E) native M^pro^ and (F) glutathionylated M^pro^ that were run as shown in (D). Shown above each peak are the molecular mass (top number) and the abundance (bottom number) found by protein deconvolution. The earlier eluting peak at 8.5 min is cm-BSA, which was used as a carrier in the runs of M^pro^ to help prevent potential non-specific losses of protein during the run. (G,H) Equilibrium analytical ultracentrifugation of (G) M^pro^ and (H) glutathionylated M^pro^ at 0.63 mg ml^-1^ (18 *µ*M) in 50 mM tris buffer pH 7.5, 2 mM EDTA, and 100 mM NaCl. The absorbance gradients in the centrifuge cell after the sedimentation equilibrium was attained at 21,000 rpm are shown in the lower panels. The open circles represent the experimental values, and the solid lines represent the results of fitting to a single ideal species. The best fit for the data shown in (G) yielded a relative molecular weight (*Mr*) of 62,991 +/− 1144 and a K_d_ for dimerization of 2.4 *µ*M and that shown in (H) yielded a molecular weight of 37,000 +/− 1000 and a K_d_ for dimerization of 200 *µ*M. The corresponding upper panels show the differences in the fitted and experimental values as a function of radial position (residuals). The residuals of these fits were random, indicating that the single species model is appropriate for the analyses.

### Modification of a single cysteine of M^pro^ leads to inhibition of dimerization and activity

To determine if glutathionylation of a single cysteine might render the enzyme monomeric and inactive, we generated a glutathionylated M^pro^ preparation by exposing 1.2 µM M^pro^ to 5 mM GSSG at pH 6.8, a pH that would favor the glutathionylation of only the most reactive cysteines (with low pKa’s). This monoglutathionylated preparation was run on SEC at 8 µM and ran as two peaks indicating the existence of both dimeric and monomeric forms of M^pro^ (Figure 4A). Deconvolution of these two peaks revealed both native and monoglutathionylated M^pro^ as expected and contained an estimated 35% monoglutathionylated M^pro^ and less than 5% diglutathionylated M^pro^, with the remaining M^pro^ unmodified (Figure 4B). However, while the mass of the unmodified M^pro^ was detected in both peaks since it is present in both monomeric and noncovalent dimeric forms (Figure 4C), the mass corresponding to monoglutathionylated protease was detected predominantly (>70% of the total area) in the second peak (Figure 4D). Treatment of the glutathionylated M^pro^ with reducing agent TCEP resulted in a decrease in the second monomeric peak (Figure 4E) and complete conversion to native M^pro^ (Figure 4F) with an elution profile consistent with native M^pro^ (Figure 4G). We also collected the first and second peaks eluting from SEC analysis of the monoglutathionylated preparation as seen in Figure 4A (peaks 1 and 2 labeled in Figure 4A) and tested them for M^pro^ activity. In the absence of 50 mM TCEP, the activity of the second peak was only 25% of that of the first peak (P<0.005) (Figure 4H). In the presence of TCEP, activity of the second peak increased significantly (P<0.01) while having no significant effect on the first peak (Figure 4H). These data provide strong evidence that monoglutathionylated M^pro^ behaves as a monomer, is inactive, and that these effects are reversible.

**Figure 4:**
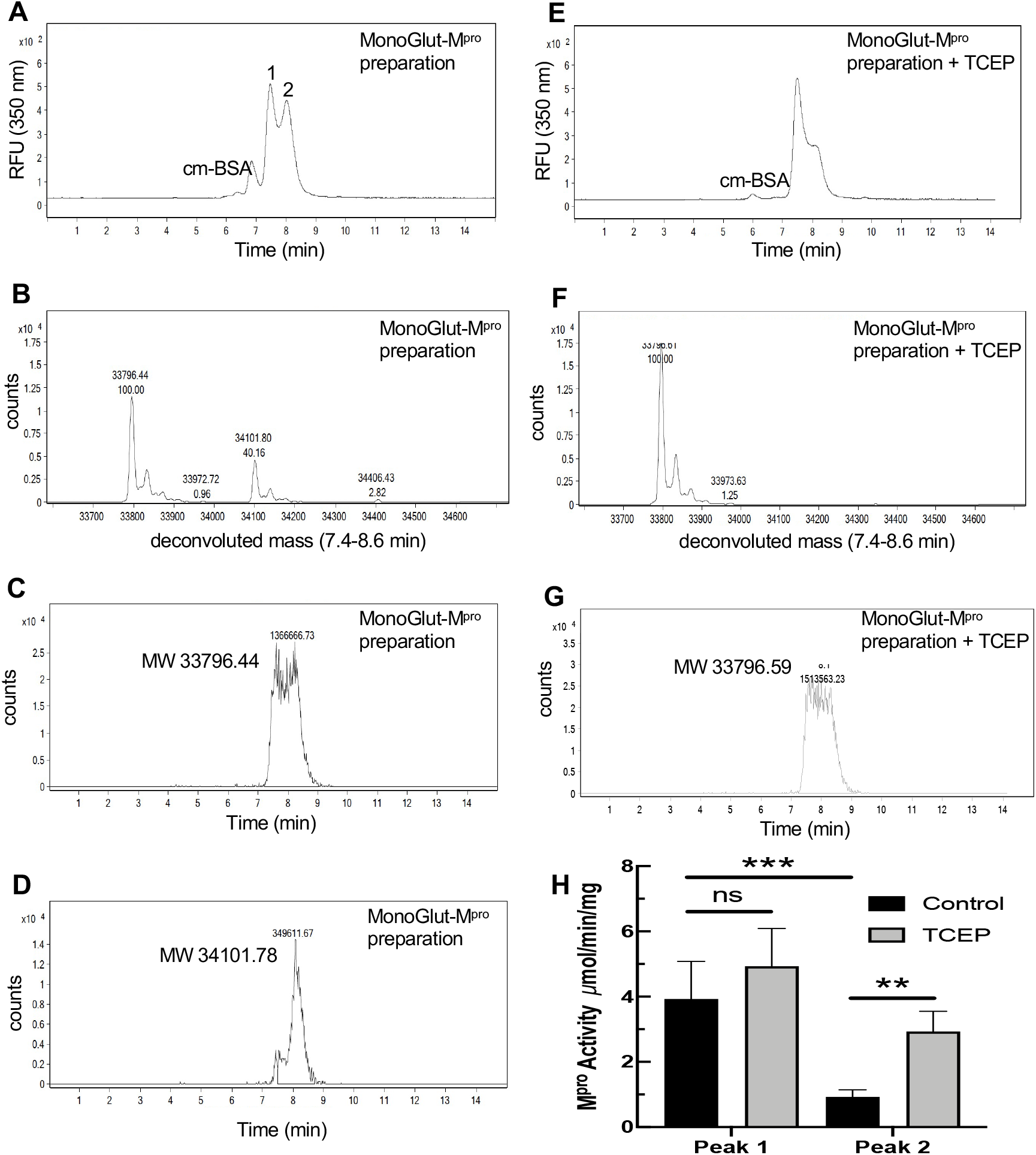
Size exclusion chromatography of a preparation of monoglutathionylated M^pro^ and analysis of M^pro^ activity. A preparation of M^pro^ containing a mixture of native and monoglutathionylated forms was made by incubating 1.2 *µ*M M^pro^ with 5 mM GSSG for 2.5 hours at 37°C, at pH 6.8, to increase specific modification of the more reactive cysteines of M^pro^ as described in Materials and Methods. (A) SEC2000 elution profile as monitored using the intrinsic protein fluorescence (excitation 276 nm, emission 350 nm) of a 2 *µ*l injection of 8 *µ*M monoglutathionylated M^pro^ preparation and (B) M^pro^ molecular weights found by protein deconvolution of the peaks in (A), (C) Elution profile for the mass of native M^pro^ in the monoglutathionylated preparation and (D) elution profile for the mass of monoglutathionylated M^pro^ in the monoglutathionylated preparation. (E) Elution profile for 2 *µ*l injection of 8 *µ*M monoglutathionylated M^pro^ preparation after treatment with 50 mM TCEP for 15 min. (F) M^pro^ molecular weights found by protein deconvolution after treatment with 50 mM TCEP for 15 min. (G) Elution profile for the mass of native M^pro^ after treatment of monoglutathionylated M^pro^ with 50 mM TCEP for 15 min. (H) M^pro^ activity without (black bars) and with (grey bars) TCEP treatment for peak #1 and Peak #2 from Fig 4A after collecting M^pro^ following SEC of the monoglutathionylated M^pro^ preparation. The values represent the average of 4 separate determinations (n=4) of M^pro^ activity. A two-way ANOVA followed by Šídak’s multiple comparison post hoc test was done. P-values less or equal to 0.05 were considered statistically significant, **<0.01 and ***<0.005 (ns= p-value > 0.05).

### Inhibition of M^pro^ activity by glutathionylation is reversible with glutaredoxin (Grx)

Grx (also known as thioltransferase) is a ubiquitous cellular enzyme that is able to reverse glutathionylation of a number of different cellular proteins including hemoglobin, nuclear factor-1, PTP1B, actin, Ras, IκB kinase, procaspase 3, and IRF-3, as well as viral proteins including HIV-1-protease and HTLV-1 protease ^19,20^. We tested whether Grx could deglutathionylate M^pro^ and restore its activity. Preparations of glutathionylated M^pro^ were prepared at pH 7.5 or pH 6.8 and then tested for reversibility of glutathionylation and restoration of activity following treatment with Grx. The glutathionylated preparation made at pH 7.5 contained no detectable unmodified M^pro^ and was predominantly diglutathionylated M^pro^ (75%) and monoglutathionylated (22%) with the remainder triglutathionylated (3%) (Figure 5A). Incubation of the preparation with 350 nm GSH alone, a cofactor required for Grx activity, produced a small amount of detectable unmodified M^pro^ (1.5%) but led to only minor changes in the percentages of the other forms of M^pro^ (Figure 5B). However, incubation of glutathionylated M^pro^ with Grx in the presence of 0.5 mM GSH resulted in the loss of the triglutathionylated M^pro^, a substantial decrease in the diglutathionylated M^pro^ (from 75% to 16%), an increase in monoglutathionylated M^pro^ (22% to 65%) and the production of native M^pro^ which made up 19% of the total M^pro^ (Figure 5C). M^pro^ activity was then assessed under these same conditions. Incubation of glutathionylated M^pro^ with 350 nM Grx in the presence of 0.5 mM GSH led to a significant increase in protease activity, restoring an average 58% of the activity compared to untreated M^pro^, while 0.5 mM GSH alone restored only about 10% of the activity (Figure 5D). We also assessed the ability of Grx to deglutathionylate the preparation made at pH 6.8. The glutathionylated preparation made at pH 6.8 contained approximately 30% monoglutathionylated M^pro^ based on percent abundance, and less than 2% diglutathionylated with the remainder (68%) being unmodified (Figure 5E). Incubation of this preparation with GSH alone for 30 min again led to insignificant changes in the percentages of monoglutathionylated M^pro^ (69.3% native, 2.9% monoglutathionylated and 1.7% diglutathionylated) (Figure 5F). However, incubation of this preparation of M^pro^ with 350 nm Grx in the presence of GSH resulted in loss of the diglutathionylated M^pro^ and a decrease in the percentage of monoglutathionylated M^pro^, going from an average 29% to 14% monoglutathionylated M^pro^ with a corresponding increase (from 69% to 86%) in unmodified M^pro^ (Figure 5G). Furthermore, Grx was found to reverse glutathionylation of M^pro^ as assessed by SEC-MALDI-TOF and restore activity in a dose dependent manner (Figure 5H), and at 175 nM, Grx restored 100% of the activity (Figure 5I). Interestingly, even at the highest concentration of Grx tested (350 nM), about 14% of the M^pro^ remained in a monoglutathionylated form (Fig 5H). This suggests that Grx is preferentially removing glutathione from cysteines whose glutathionylation is responsible for inhibition of activity while sparing certain cysteines whose modification does not alter activity.

**Figure 5:**
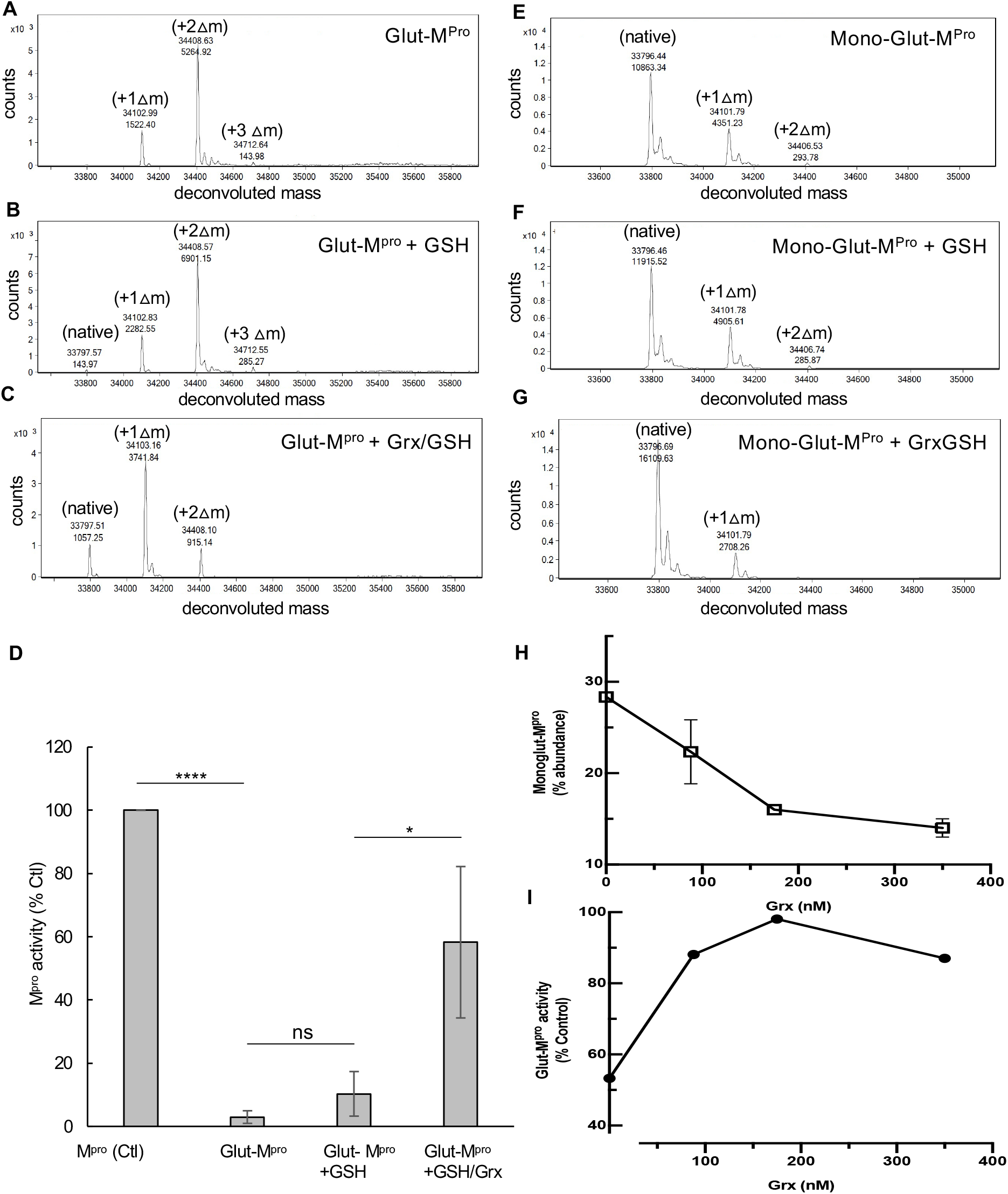
Grx reverses glutathionylation and restores M^pro^ activity. (A-C) M^pro^ Glutathionylated at pH 7.5 was incubated (3 *µ*M final) for 30 min in the presence of (A) buffer control, (B) GSH (0.5 mM) or (C) GSH (0.5 mM) with Grx (final 350 nM). Samples were analyzed by SEC3000/MALDI-TOF and the eluting protease analyzed by protein deconvolution (8.3-10 min) to determine the M^pro^ species present. The experimental masses (top number) are shown as well as the abundances (bottom number) for each peak obtained by deconvolution. The native M^pro^, as well as the increases in masses indicative of glutathionylation, are indicated for the addition of 1 (+□1), 2 (+□2), and 3 (+□3) glutathione moieties in the deconvolution profiles. (D) Samples of glutathionylated M^pro^ were treated as in (A-C) and then analyzed for M^pro^ activity and compared to unmodified M^pro^. M^pro^ activity for control in (D) was 5.77+/− 1.5 *µ*M/min/mg, and percent activity for the different conditions was normalized to their respective controls. (E-G) Monoglutathionylated M^pro^ was incubated (8 *µ*M final) for 15 min in the presence of (A) buffer control, (B) GSH (0.1 mM) or (C) GSH (0.1 mM) with Grx 350 nM and samples analyzed by SEC2000/MALDI-TOF deconvolution (7.3-8.6 min). (H, I) Samples were prepared as in (E-G) and the percentage of monoglutathionylated M^pro^ and activity was determined after the 15-minute incubation with 0, 88, 175, or 350 nm Grx in the presence of 100 *µ*M GSH. (H) Percent of monoglutathionylated Mpro after Grx treatment and (I) M^pro^ activity after Grx treatment. The M^pro^ activity was normalized to the TCEP treated preparation which yielded fully reduced native M^pro^ and was used as 100% activity. For (D) Values represent the average +/− standard deviation of 4 separate experiments (* = p-value < 0.05, ****=p-value < 0.001 paired students t-test, ns=not significant p>0.05). For (H) the values are the average of 3 separate experiments (n=3) and for (I) one experiment performed in duplicate (n=2).

### Identification of glutathionylated cysteines by MALDI-TOF MS

To determine which cysteines of M^pro^ might be primarily responsible for the inhibition of dimerization and activity, we digested native M^pro^ and a monoglutathionylated preparation of M^pro^ (containing approximately 35% monoglutathionylated forms of M^pro^) with either chymotrypsin or a combination of trypsin and lysC to produce peptides that could be assessed for glutathionylation. Prior to digestion, we alkylated the free cysteines in the M^pro^ preparations with N-ethylmaleimide (NEM) using the AccuMAP™ System (Promega); this step limits disulfide scrambling during the alkylation and proteolytic digestion processes. For digestions of native M^pro^ (see Figure S2A for TIC chromatogram and S2B for UV chromatogram in supplemental material) that was fully alkylated with NEM, we were able to identify alkylated peptides for 7 of the 12 cysteines of M^pro^ including cysteines 38, 44, 117, 128, 145, 156 and 300 by using molecular ion extraction for the predicted monoisotopic masses (see peptides 1-10 in Table S1 in supplemental material) along with 12 other non-cysteine containing peptides (see peptides 15-27 in Table S1 in supplemental material). To identify which cysteines were becoming glutathionylated in the glutathionylated M^pro^ preparation (see Figure S2C for TIC chromatogram and S2D for UV chromatogram in supplemental material), we searched for the predicted glutathionylated monoisotopic masses by molecular ion extraction of the TIC chromatogram obtained from RP-HPLC/MALDI-TOF analysis of chymotrypsin digests. We located monoisotopic masses consistent with that for three glutathionylated peptides (glutathione adds a net 305.08 amu): peptides ^151^NIDYDC^GSH^VSF^159, 295^DVVRQC^GSH^SGVTF^305^ and ^295^DVVRQC^GSH^SGVTFQ^306^ with glutathionylated Cys^156,^ Cys^300^, and Cys^300^ respectively (Table 1 and see Figure S3A-S3J for detailed analysis in supplemental material). All three of these peptides had experimental masses that were within 0.04 amu of the predicted glutathionylated masses (predicted monoisotopic mass increase with glutathione is 305.08) consistent with addition of glutathione. To confirm that these peptides were, indeed, glutathionylated forms of the predicted M^pro^ peptides, we analyzed the peptide digests both before and after treating them with TCEP to reduce any disulfide bonds (see Figure S2E for TIC chromatograms and S2F for UV chromatograms in supplemental material). When this was done, the masses for all three of the predicted glutathionylated peptides were no longer detected, due to the removal of glutathione with TCEP, and in place we were able to locate the predicted native masses expected following removal of glutathione for all three peptides (Table 1 and see Figure S3K-S3P in supplemental material). The difference (Delta) between the experimental and calculated masses was less than 0.05 amu for all peptides providing strong confidence in their identity (Table 1).

**Table 1:**
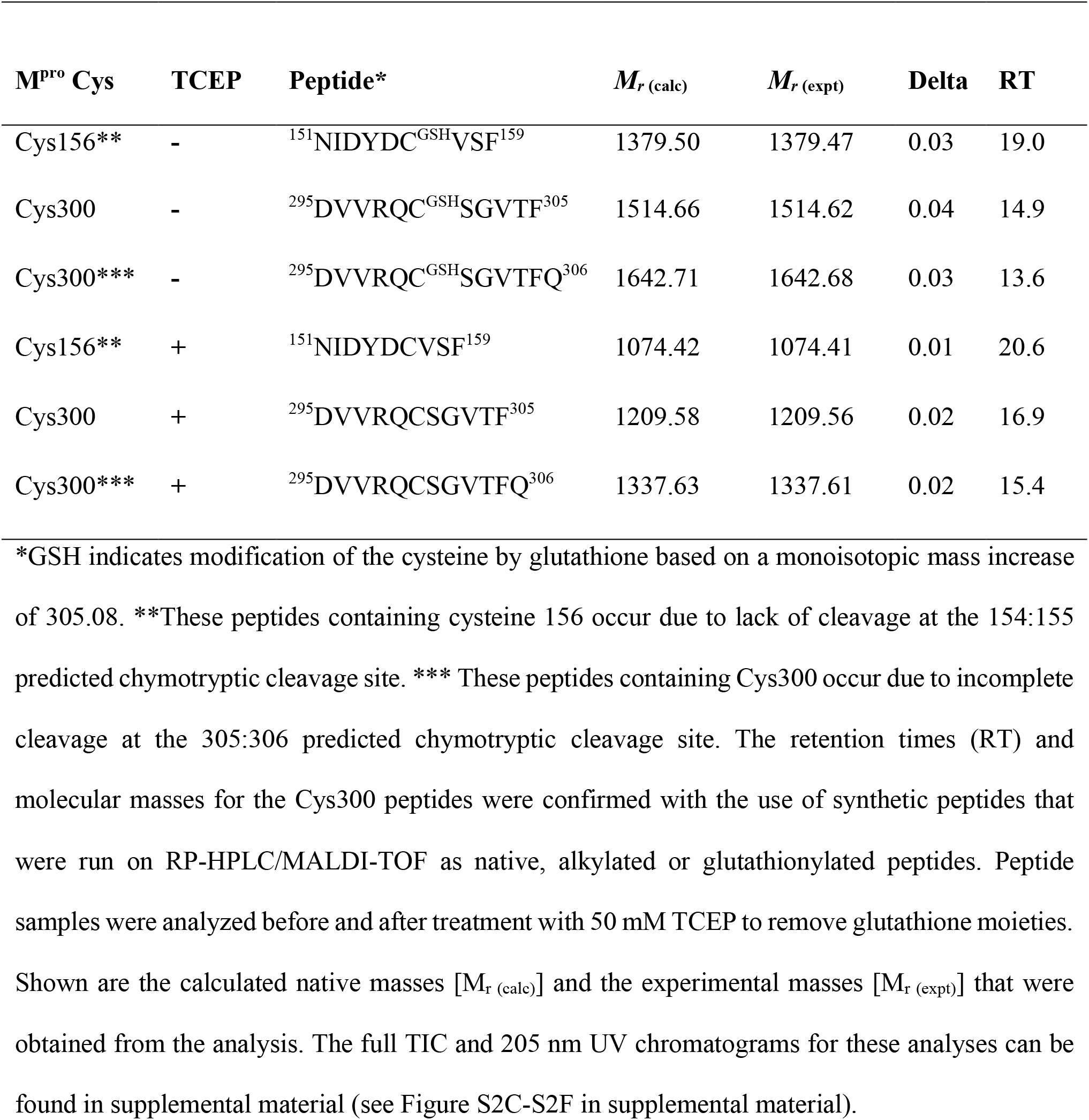
RP/HPLC/MALDI-TOF MS Identification of peptides from chymotrypsin digestion of monoglutathionylated M^pro^ preparations without (-) and with (+) TCEP.

Due to the inability to assess modification of cysteines 16, 22, 85, 161 and 265 using the chymotrypsin data, as the peptides carrying these residues were not located (see Table S1 for a list of the peptides found in supplemental material), we prepared trypsin/lysC digests of native M^pro^ and the same monoglutathionylated M^pro^ preparation used in the chymotrypsin experiments (see Figure S4A,C for TIC chromatogram and S5B,D for UV chromatograms). Interrogation of the TIC chromatogram for masses corresponding to glutathionylated forms of cysteine-containing peptides revealed masses consistent with glutathionylation of three peptides: ^77^VIGHSMQNC^GSH^VLK^88, 299^QC^GSH^SGVTFQ^306^ and ^299^pyQC^GSH^SGVTFQ^306^ (the pyroglutamate (py) form of the 299-306 peptide that results from spontaneous deamidation of peptides with N-terminal glutamyl residues^21^) (Table 2 and see Figure S5A-S5J in supplemental material). These were glutathionylated at Cys^85^, Cys^300^, and Cys^300^, respectively. All three peptides had experimental masses within 0.04 amu of the predicted calculated glutathionylated masses consistent with glutathione modification (Table 2). Also, the calculated masses for the three native forms were found following analysis of the tryptic digests after reduction with TCEP (Table 2 and see Figure S5E-S5P in supplemental material). The difference (Delta) between the experimental and calculated masses was less than 0.05 amu providing strong confidence in their identity (Table 2). The data from the trypsin/lysC digestion indicated that the majority of the monoglutathionylation was occurring at Cys300. We based this on the greater area at 205 nm obtained for glutathionylated Cys300 peptides than the cys85 peptide (combined area for glutathionylated cys300 peptides at 205 nm was 301 vs 56 for the glutathionylated cys85 peptide) and their native forms (combined area at 205 nm for native cys300 peptides was 272 vs 21 for the native cys85 peptide) (see Figure S5C-S5D in supplemental material). Taken together, the data obtained from the chymotryptic and tryptic/lysC digestions of M^pro^ and glutathionylated M^pro^ strongly implicated Cys300 as a primary target for glutathionylation. Given the location of Cys300 near the dimer interface and the importance of amino acids 298 and 299 for dimerization ^4,7^ we hypothesized that glutathionylation of this cysteine is likely responsible for interfering with dimerization leading to inhibition of M^pro^ activity.

**Table 2:** RP/HPLC/MALDI-TOF MS Identification of peptides from trypsin/lysC digestion of monoglutathionylated M^pro^ preparations without (-) and with (+) TCEP.

| M<sup>pro</sup> Cys | TCEP | Peptide* | $M_r(\text{calc})$ | $M_r(\text{expt})$ | Delta | RT |
| --- | --- | --- | --- | --- | --- | --- |
| Cys85 | - | <sup>77</sup>VIGHSMQNC<sup>GSH</sup>VLK<sup>88</sup> | 1632.74 | 1632.71 | 0.03 | 13.5 |
| Cys300 | - | <sup>299</sup>QC<sup>GSH</sup>SGVTFQ<sup>306</sup> | 1173.44 | 1173.42 | 0.02 | 10.9 |
| Cys300** | - | <sup>299</sup>pyQC<sup>GSH</sup>SGVTFQ<sup>306</sup> | 1156.44 | 1156.40 | 0.04 | 13.6 |
| Cys85 | + | <sup>77</sup>VIGHSMQNCVLK<sup>88</sup> | 1327.66 | 1327.64 | 0.02 | 14.7 |
| Cys300 | + | <sup>299</sup>QCSGVTFQ<sup>306</sup> | 868.36 | 868.36 | 0.00 | 11.2 |
| Cys300** | + | <sup>299</sup>pyQCSGVTFQ<sup>306</sup> | 851.36 | 851.33 | 0.03 | 14 |

\*GSH indicates modification by glutathione based on a monoisotopic mass increase of 305.08.
\*\*These peptides are the result of the spontaneous deamidation that occurs with peptides containing an N-terminal glutamyl residues <sup>21</sup> and the retention times and molecular masses for this peptide were confirmed with the use of synthetic peptides that were run on RP-HPLC/MS. The retention times (RT) and molecular masses for the Cys300 peptides were confirmed with the use of synthetic peptides that were run on RP-HPLC/MALDI-TOF as native, alkylated or glutathionylated peptides. Peptide samples were analyzed without (-) and with (+) TCEP to remove glutathione moieties. Shown are the calculated native masses [ $M_r(\text{calc})$ ] and the experimental masses [ $M_r(\text{expt})$ ]. The full TIC and 205 nm UV chromatograms for these analyses can be found in supplemental material (see Figure S5C-S5F in supplemental material).

### Cys300 is required for inhibition of M^pro^ activity following glutathionylation

To determine if Cys300 was contributing to the inhibition of activity of M^pro^ following glutathionylation, we prepared a C300S mutant M^pro^ (for purity and molecular weight analysis see Figure S1F-S1I) and evaluated the effects of glutathionylation on M^pro^ activity. We treated WT and C300S M^pro^ at 1.2 µM with 10 mM GSSG for 30 minutes and then measured activity. In these experiments, the activity of WT M^pro^ was inhibited by more than 50% while C300S M^pro^ was not significantly affected (Figure 6A). We also measured the extent of glutathionylation for WT and C300S M^pro^ following the enzyme assay. Based on the absolute abundances of each form, we found that WT M^pro^ had 46%, 14% and 5% mono, di and triglutathionylated forms, respectively, with the remainder (35%) unmodified while after the same treatment, C300S had 36% and 11% mono and diglutathionylated forms, respectively, with the remainder (53%) unmodified (see Figure S6A-S6D in supplemental material). This indicated that while almost 50% of C300S could still become glutathionylated at other cysteine residues, its activity was unaffected, strongly implicating Cys300 in the inhibition of M^pro^ activity following glutathionylation of WT M^pro^. To determine if Cys300 was the primary target for glutathionylation when incubating with GSSG at the lower pH of 6.8, we treated WT and C300S M^pro^ with 5 mM GSSG at pH 6.8 for 2.5 hours to produce monoglutathionylated forms of M^pro^. Based on SEC/MALDI-TOF analysis the WT M^pro^ was 36% glutathionylated while the C300S M^pro^ was only 16% glutathionylated based on the abundances for each form (supplemental Figure S6E-S6F). This data suggests that there are at least two reactive cysteines under these lower pH conditions. Activity of these preparations was measured before and after reduction with DTT. DTT increased the activity of the monoglutathionylated WT M^pro^ preparation by 26% but had no significant effect on the activity of monoglutathionylated C300S M^pro^ mutant (Figure 6B). This suggests that while the C300S mutant can still become glutathionylated at alternative cysteines, the modification has little effect on M^pro^ activity.

**Figure 6:**
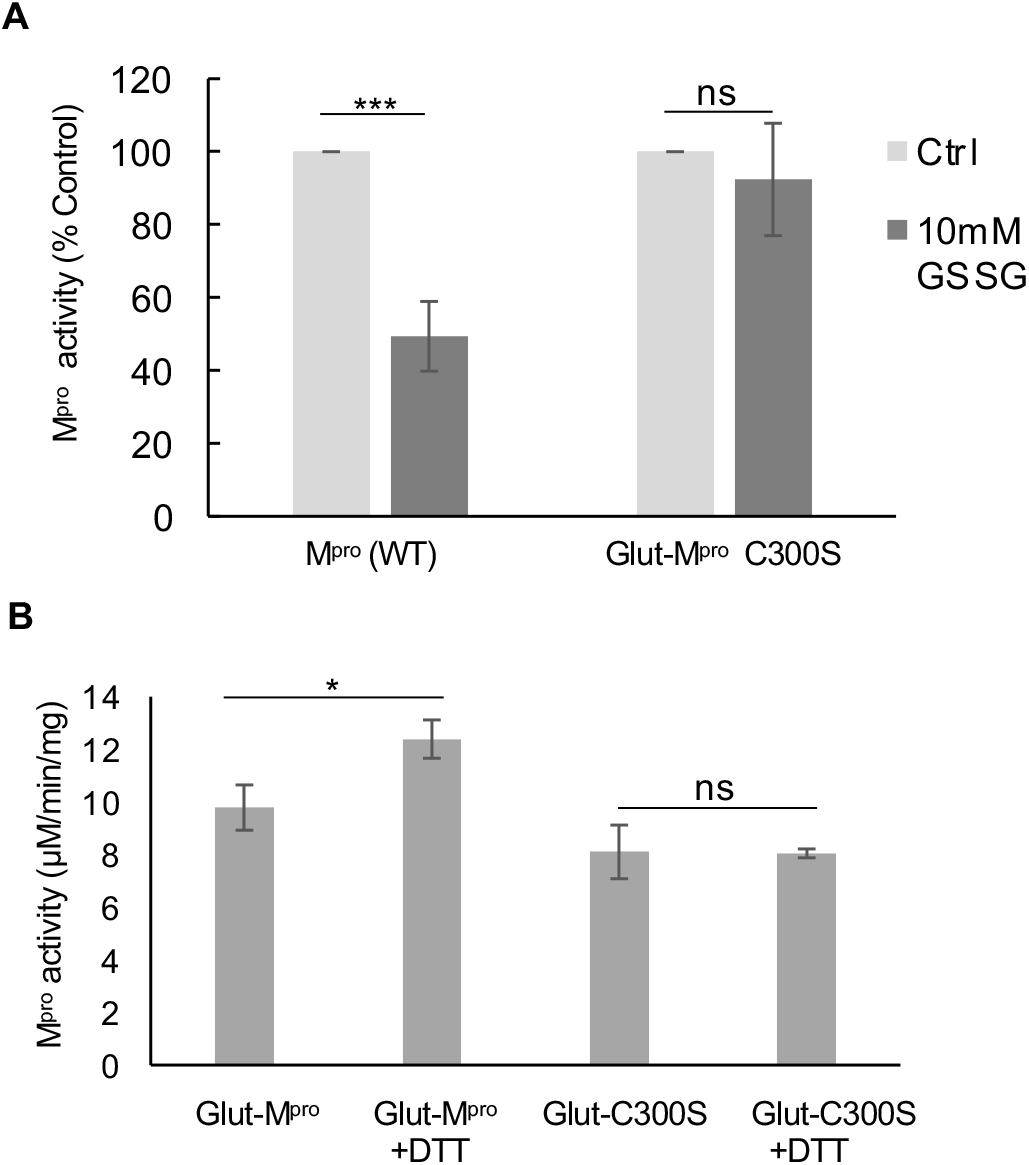
Glutathionylation inhibits WT SARS-Cov-2 M^pro^ activity but not C300S M^pro^ activity. (A) Activity of wild type (WT) and C300S M^pro^ (1 *µ*M enzyme) following a 30-minute pre-incubation of 1.2 *µ*M M^pro^ with 10 mM oxidized glutathione. (B) M^pro^ activity for a WT monoglutathionylated M^pro^ preparation (containing approximately 30% monoglutathionylated M^pro^ and 4% diglutathionylated) and a C300S monoglutathionylated M^pro^ preparation (containing approximately 18% monoglutathionylated M^pro^) preincubated for 10 minutes without or with 20 mM DTT. The amount of monoglutathionylated M^pro^ was estimated using the relative abundances of native M^pro^ and glutathionylated M^pro^ following deconvolution of the eluting M^pro^ species from SEC/MALDI-TOF analysis. Values represent the average +/− standard deviation of 3 separate experiments (n=3) (* = p-value < 0.05, ***=p-value < 0.005 paired students t-test, ns=not significant p>0.05).

## Discussion

In cells that are under oxidative stress, cellular and foreign proteins can undergo glutathionylation, and this process, which is reversible, can alter the function of these proteins ^18,19,22,23,24^. Biochemical studies with GSSG can be carried out to determine if reversible glutathionylation might regulate the activity of key proteins although glutathionylation of proteins within cells more likely goes through sulfenic acid intermediates^19^. In this study, we show that glutathionylation of SARS-CoV-2 M^pro^ inhibits M^pro^ activity, and this is reversible with reducing agents or the ubiquitous cellular enzyme, Grx. We also show that loss of activity is due to inhibition of M^pro^ dimerization following modification of Cys300. Cys300 of M^pro^ is located proximal to Arg298 and Gln299, both of which play pivotal roles in M^pro^ dimerization in the C-terminal dimerization domain ^4^. Our data indicate Cys300 is particularly sensitive to glutathionylation, as we were able to modify Cys300 at pH 6.8, a pH where cysteines are usually protonated and unreactive due to typical pKa’s around pH 8. Our current model for regulation of dimerization and activity of M^pro^ is shown in Figure 7A. Our data indicates that monomeric M^pro^ is susceptible to glutathionylation at Cys300 and this blocks dimerization. Grx can reverse the modification, thus restoring dimerization and activity of M^pro^ (Figure 7A). We hypothesize that glutathionylation of Cys300 in SARS-CoV-2 infected cells would inhibit M^pro^ activity and therefore decrease SARS-CoV-2 replication during oxidative stress. Thus, SARS-CoV-2 M^pro^, and by analogy SARS-CoV-1, are quite similar to retroviral proteases in being essential for viral replication, requiring dimerization for activity, and being susceptible to reversible inhibition by glutathionylation ^8, 9, 11, 13, 14, 15, 16^.

**Figure 7:**
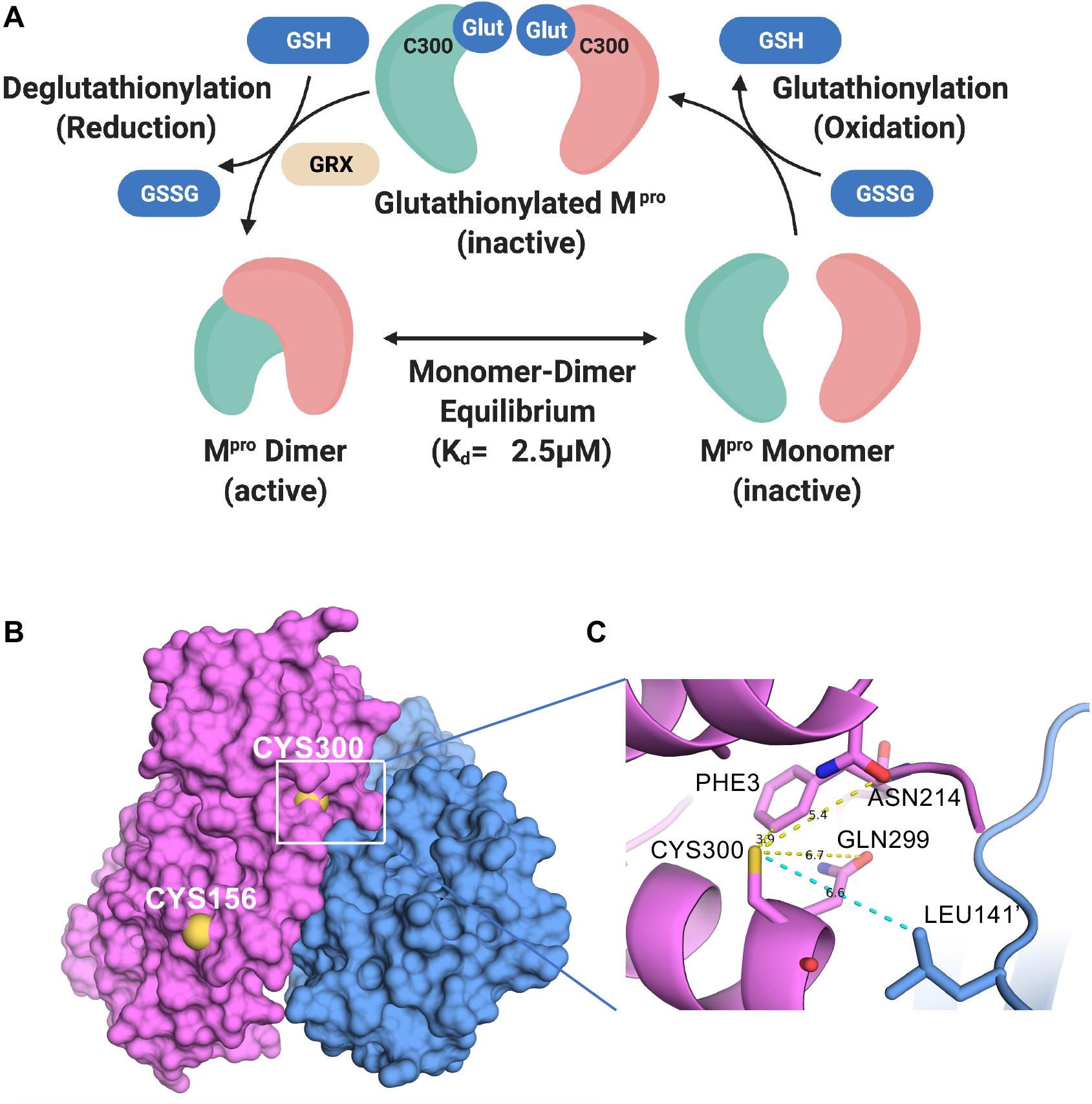
The current model for the regulation of dimerization and activity through reversible glutathionylation of M^pro^ and Space filling and close up ribbon model of SARS-CoV-2 M^pro^. (A) Model showing that M^pro^ dimer exists in equilibrium with its monomer form with a determine K_d_ of 2.5 *µ*M. The monomeric M^pro^ is susceptible to glutathionylation at Cys300, and this leads to inhibition of dimerization and loss of activity. Human Grx is able to reverse glutathionylation of Cys300 and restore dimerization and activity. (B) Space filling model of the SARS-CoV-2 M^pro^ dimer (apo form) showing the location of cysteines 156 on the surface and 300 near the dimer interface in the left (pink) protomer (PDB ID 7K3T). (C) Close up ribbon model around Cys300 showing the proximity to protomer 2 (blue) at leucine 141’ and the proximity to ASN214, GLN299 and PHE3.

Identification of which cysteines in SARS-CoV-2 M^pro^ are glutathionylated was not a trivial matter as M^pro^ contains 12 cysteine residues all in their reduced form. For this reason, we used the Accumap™ low pH system to alkylate M^pro^ with NEM to minimize disulfide scrambling during the reactions. Our studies indicated that at least two cysteines were readily modified by GSSG including Cys300 and Cys156 (Figure 7B). We identified glutathionylated peptides by their predicted monoisotopic masses and the alkylated forms of these peptides in controls using RP/HPLC/MALDI-TOF, and also showed the disappearance of these masses after reduction with TCEP leading to the appearance of their native peptide counterparts. The identity of Cys300 glutathionylated and native and alkylated peptides were further confirmed with the use of synthetic peptides used as standards to determine masses and retention times. The data from chymotryptic and tryptic/lysC digestions implicated Cys300; therefore, we prepared a C300S M^pro^ mutant to verify the role of cysteine Cys300 in inhibition by glutathionylation. Indeed, C300S M^pro^ was no longer susceptible to inhibition by glutathionylation under the same conditions where WT M^pro^ was, thus confirming the role for Cys300 in this process.

Glutathionylation of proteins occurs via a mixed disulfide between glutathione and a cysteine residue. Most cysteine residues have relatively high pKa’s (pH 8.0 or greater) and usually remain protonated under physiologic conditions, making them relatively unreactive at typical cellular pH. However, studies have shown that the local environment around certain cysteine residues can lower their pKa making them more susceptible to oxidation and glutathionylation ^25,26,27^. We propose that the local environment of Cys300 may account for this particular susceptibility to glutathionylation. Previous studies have found that the presence of basic residues or serine hydroxyl sidechains in the local environment can substantially reduce the pKa of the thiol sidechain ^25,28^. As to Cys300, there is a basic residue at Arg298 and a hydroxyl residue at Ser301. This may increase the local acidity of the Cys300 thiol group in the monomer making it more prone to oxidation while in the dimeric state Arg298 is involved in interactions which stabilize the dimer ^7^. In the SARS-CoV-2 dimer Inspection of a previously determined monomeric form of SARS-CoV-1 M^pro^ (R298A) reveals that the carbonyl sidechains of Asn214 and Gln299, which can act as hydrogen acceptors and potentially destabilize the thiol group, have close contact with the Cys300 thiol (Figure 8). Although there is not a monomer structure of SARS-CoV-2 M^pro^ the distances of the Cys300 thiol to the carbonyls in SARS-CoV-1 and 2 is much greater, possibly decreasing its reactivity (see Figure S7A and S7B in supplemental material).

**Figure 8:**
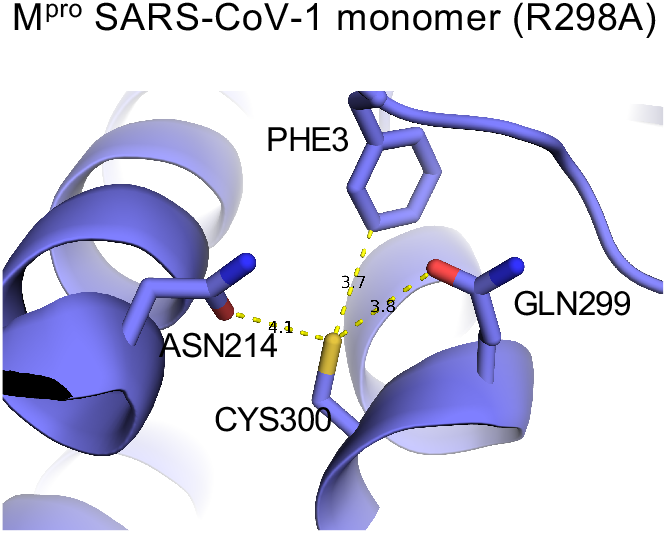
The local environment around Cys300 in monomeric SARS-CoV-1 M^pro^. Ball and stick model for local environment around cys300 in R298A M^pro^ monomer PDB ID 2QCY (a monomeric form of SARS-CoV M^pro^ mutant R298A at pH 6.0). Structural figures were produced with PyMOL v1.5.0.4 ^40^.

It is possible that regulation of M^pro^ through reversible oxidation/glutathionylation of M^pro^ may have evolved in part as a mechanism to blunt viral processing and replication in cells undergoing significant oxidative stress which otherwise may generate defective viral particles. It’s known that viral infection itself leads to oxidative stress in cells even early on in infection ^29^. In the case with SARS-CoV-2, Cys300 may act as a sensor to regulate when viral proteolytic processing should take place to optimize the generation of new virions. Moreover, M^pro^ from SARS-CoV-1 and SARS-CoV-2 contain 12 cysteines and 10 methionine residues. Studies have shown that such residues can act as decoys to prevent permanent damage to proteins during oxidative stress ^30,31^. In the case of M^pro^, this could help protect the active site cysteine required for catalysis. It should be noted that the details of the initial autocatalytic processing of M^pro^ from the polyprotein pp1a and pp1ab are still not fully understood, but in the case of HIV, we have shown that similar modifications can also affect the initial autocleavage of the Gag-Pol-Pro polyprotein ^11,16^.

Another possible factor that may have led to this feature of coronavirus M^pro^ relates to its evolution in bats. It’s important to point out that the M^pro^’s from the three closest relatives to SARS-CoV-2 derived from bats ^32^ have an extremely high degree of amino acid identity (see Figure S8 in supplemental material) to that of SARS-CoV-2 and all three contain 12 cysteine residues including Cys300. SARS-CoV-2 is thought to have jumped to humans from an original reservoir in *Rhinolophus* bats, possibly through an intermediate host ^33^. Bats are reservoirs for a vast number of coronaviruses and other RNA viruses and are often infected with these viruses without showing any signs of disease ^34^. One reason for this coexistence is that bats have evolved an immune response to RNA viruses with substantial interferon activity but a minimal inflammatory response ^34^. The act of flying requires considerable metabolic energy, and when in flight and during migration, bats are placed under high levels of oxidative stress ^35,36,37^. Moreover, bats spend much of their lives in densely populated shelters such as caves that facilitate virus transmission. Maintaining the health of host bat colonies would appear to be a good evolutionary strategy for coronaviruses and one can speculate that SARS-CoV-1 and SARS-CoV-2 and related RNA bat viruses have co-evolved so as to persist in bat colonies by not killing off their host animals. Part of this evolutionary adaption might be dampening of viral replication under conditions of oxidative stress, through the inhibition of M^pro^ by glutathionylation. At this time, it is unclear what ramifications these effects from M^pro^ glutathionylation might have for SARS-CoV-2 infection in humans. Unlike bats, humans are not exposed to the metabolic and oxidative stress that is encountered in bats during flight and therefore would not be expected to suppress SARS-CoV-2 replication through this mechanism. This may help explain the relatively more severe manifestations of SARS-CoV-2 infection in humans than in bats.

A more practical implication of our findings is that it can inform the development of anti-viral drugs against SARS-CoV-2. While vaccines are effective at preventing COVID-19, effective anti-SARS-CoV-2 drugs are urgently needed and will be in the foreseeable future. Because of its essential role in SARS-CoV-2 replication, M^pro^ is an attractive target for drug development. Nearly all of this effort has focused on active site inhibitors of M^pro^ which can block SARS-CoV-2 replication and cytopathic effect ^1,2,6 38^. Our observation that Cys300 at the dimer interface is particularly susceptible to oxidative modification, and that this modification can block dimerization of M^pro^ resulting in inhibition of activity, demonstrates an alternative way of targeting M^pro^. Being on the M^pro^ surface in the monomer, this cysteine may be highly accessible and may thus be a promising target for the development of specific M^pro^ inhibitors. In this regard, Gunther and Reinke et al.^38^ have recently identified the hydrophobic pocket consisting of Ile21, Leu253, Gln256, Val297, and Cys300 of SARS-Cov-2 Mpro as an allosteric binding site for non-active site M^pro^ inhibitors. Our results indicate that this area can be specifically targeted through Cys300, which is highly reactive and leads to inhibition of dimerization.

## Materials and Methods

### Enzymes, peptides and reagents

The substrate peptide for M^pro^ (H2N-TSAVLQ-pNA) and peptides corresponding to some of the predicted chymotryptic fragments containing cysteine residues including M^pro^ peptide fragments 113:118, 127:134, 141:150, 155:159, 295: 305 and 295: 306 as well the predicted tryptic fragment, 299:306, were obtained (>95% purity) from New England Peptide (Gardner, MA). Amicon Ultra-Centrifugal Filters (10 kDa cutoff, 0.5 ml and 15 ml), carboxymethyl bovine serum albumin (cm-BSA), oxidized and reduced forms of L-glutathione (Bioxtra) (>98%), 4-nitroaniline (>99%), the reducing agents Tris (2-carboxyethyl) phosphine hydrochloride (TCEP) and dithiothreitol (DTT) were from Sigma-Aldrich (Milwaukee, WI). BioSep SEC3000 and SEC2000 size exclusion columns (300 × 4.6 mm) were from Phenomenex (Torrence, CA). The VydacC18 column (218TP5205) was from MAC-MOD Analytical (Chadds Ford, PA). Peptide desalting columns from ThermoFisher Scientific (Pittsburgh, PA) and AccuMap™ low pH protein digestion kit (with trypsin and lysC) and chymotrypsin (sequencing grade) were from Promega (Madison, WI). PreScisson protease was from GenScript (Piscataway, NJ). Recombinant human glutaredoxin (Grx) transcript variant 1 was from Origene (cat# TP319385) (Rockville, MD) and stored at −70°C in 25 mM Tris.-HCl, pH 7.3, 100 mM glycine and 10% glycerol (7 µM stock).

### Expression and purification of Authentic M^pro^ and C300S M^pro^

The SARS-CoV2 M^pro^-encoding sequence and C300S mutant sequence were cloned into pGEX-4T1 vector (Genscript) with N-terminal self-cleavage site (SAVLQ/SGFRK) and C-terminal His_6_-tag as previously designed by others ^6^. The plasmid constructs were transformed into BL21 Star™ (DE3) cells (Thermo Fisher Scientific). The cultures were grown in Terrific Broth media supplemented with ampicillin (Quality Biological, Gaithersburg, MD). Protein expression was induced by adding 1 mM iso-propyl b-D-thiogalactopyranoside at an optical density of 0.8 at 600 nm and the cultures were maintained at 20 °C overnight. SARS-CoV2 M^pro^ and C300S M^pro^ were purified first by affinity chromatography using TALON™ cobalt-based affinity Resin (Takara Bio). The His_6_-tag was cleaved off by PreScission protease and the resulting authentic 306 amino acid M^pro^ (see Figure S1A in supplemental material) and C300S M^pro^ were further purified by SEC using a HiLoad Superdex 200 pg column (GE Healthcare) in 20 mM Tris, pH 7.5, 150 mM NaCl, and 2 mM DTT. The purity and molecular mass of M^pro^ were assessed by LDS-gel electrophoresis as well as reverse phase high performance liquid chromatography (RP/HPLC) on a C18 column coupled with a Matrix-Assisted Laser Desorption/Ionization-Time of Flight (MALDI-TOF) mass spectrometer (MS). The purity of these M^pro’^s was greater than 95% by LDS-gel electrophoresis, RP-HPLC chromatography (205 nm), and MALDI-TOF analysis (see Figure S1B-S1E in supplemental material), with an average experimental mass of 33796 amu +/− 1 amu (expected average mass of 33796.48 amu) (see Figure S1E and S1I (insets) in supplemental material). Final preparations of M^pro^ (2-6 mg/ml) were stored at −70 in 40 mM Tris-HCl buffer, pH 7.5, 2 mM DTT and 150 mM NaCl.

### M^pro^ colorimetric enzyme assay

The enzymatic activity of M^pro^ of SARS-CoV-2 was measured using the custom-synthesized peptide, H2N-TSAVLQ-pNA as described previously ^39,40^. TSAVLQ represents the nsp4↓nsp5 cleavage sequence for SARS and SAS2 M^pro^. The rate of enzymatic activity was determined by following the increase in absorbance (390 nm) using a Spectramax 190 multiplate reader at 37°C as a function of time following addition of substrate. Assays were conducted in clear flat bottom 96-well plates (Corning) containing 40 µL of assay buffer (50 mm Tris, pH 7.5, 2 mM EDTA, and 300 mM NaCl containing 100 ug/ml of cm-BSA). Reactions were started by the addition of 10 µl of 2 mM substrate dissolved in ultrapure water. Activity was obtained by measuring the increase in absorbance at 390 nm as a function of time within the linear range of the assay. A calibration curve was obtained for the product, 4-Nitroanaline (pNA), and was used to convert the rate of the reaction to units of micromoles of product per min per mg of protein(μm/min/mg). In some cases, activity and M^pro^ modifications were determined by first stopping the assay at a set time by acidification with formic acid (FA)/trifluoroacetic acid (TFA) and then analyzed by RP-HPLC using a 2% acetonitrile gradient on a Vydac C18 column as described below. The activity was calculated based on the amount of pNA product (detected at 390 nm). Unprocessed substrate with detected at 320 nm.

### Glutathionylation of M^pro^ at pH 7.5 and pH 6.8

To prepare glutathionylated M^pro^ for use in analytical ultracentrifugation, SEC and activity assays, M^pro^ was first exchanged into a buffer containing 40 mM tris-HCL, 2 mM EDTA and 300 mM NaCl at Ph 7.5 using Amicon 10 kDa cutoff filter units. M^pro^ (1.2-2.2µM as noted in the Results) was then treated only with buffer or with a final of 10 mM GSSG diluted from a stock of 200 mM GSSG that had been adjusted to neutral pH with sodium hydroxide. The solutions were then incubated at 37°C for 60 min or otherwise as described in the results before removing excess GSSG. Preparations were then diluted 10X with buffer (50 mM tris-HCL, 2 mM EDTA and 100 mM NaCl) and washed 4 times using Amicon 10 kDa cutoff filter units (0.5 ml) to remove excess GSSG. The final preparations were concentrated further with a 0.5 ml 10 kDa filtration unit (0.6 mg/ml). In some cases, these preparations were concentrated to 2-6 mg/ml) for use in SEC. While the extent of glutathionylation varied among preparations the procedure usually yielded preparations of M^pro^ that contained predominantly diglutathionylated M^pro^ based on MS deconvolution analysis as well as monoglutathionylated and triglutathionylated forms.

To selectively modify M^pro^ with GSSG on the more reactive cysteine residues, a similar procedure to that above was used except 5 mM GSSG was used and we lowered the buffer pH to 6.8. Prior to modification, M^pro^ was treated with 50 mM TCEP for 30 minutes to ensure all cysteines were in their reduced form and then TCEP removed by multiple washes through an Amicon 10 kDa cutoff filter with pH 6.8 incubation buffer (50 mM tris-HCL, 2 mM EDTA and 100 mM NaCl). For glutathionylation, M^pro^ (1.2 µM) was incubated for 2.5 hours at 37°C in 50 mM Tris-HCl buffer, 300 mM NaCl, and 2 mM EDTA at pH 6.8 with buffer (control) or 5 mM GSSG. The preparations were then washed 4 times to remove excess GSSG using Amicon 10 kDa cutoff filter units (0.5 ml) with pH 6.8 buffer. This procedure typically resulted in 30-40% of becoming monoglutathionylated with less than 10% diglutathionylated. The percent of the glutathionylated M^pro^ forms was estimated based on the abundances of the different protein forms (obtained by protein deconvolution). Although these forms are similar in molecular weight, they would have somewhat different ionization potentials and therefore the numbers are only an estimate of percent modification.

To confirm the identity of certain peptide fragments we purchased synthetic peptides and modified them accordingly and determined their masses and retention times on the RP-HPLC/MS analysis. Peptides (100 µM) corresponding to chymotryptic fragments from digested M^pro^ (113:118, 127:134, 141:150, 155:159, 295: 305) were glutathionylated with 10 mM GSSG in 50 mM Tris-HCl buffer, 300 mM NaCl, and 2 mM EDTA pH 7.5 for 1 hour. These same peptides as well as 295: 306 and the tryptic peptide 299:306 were alkylated with 5 mM NEM for 30 minutes at 37 °C then acidified to pH less than 3.0 with formic acid. Glutathionylation and NEM alkylation of the peptides was verified using RP-HPLC/MS TOF analysis on a Vydac C18 column with the same method that was used for analysis of trypsin/lysC and chymotrypsin digests of M^pro^ as described below.

### Grx Assays on Glutathionylated forms of M^pro^

To determine if Grx could deglutathionylate M^pro^, monoglutathionylated preparations of M^pro^ containing 30-40% monoglutathionylated or multiglutathionylated M^pro^ (prepared as described in “Glutathionylation of M^pro^ at pH 7.5 and pH 6.8”) (8 µM) were used. For preparations made at pH 7.5 which had predominantly diglutathionylated M^pro^ the preparation was incubated at 37°C for 30 minutes in the presence of buffer control (50 mM Tris, pH 7.5, 2 mM EDTA, and 100 mM NaCl containing 100 ug/ml of cm-BSA), Grx (350 nM) alone, GSH alone (0.5 mM) and Grx and GSH together. The samples were then analyzed for M^pro^ activity and by SEC3000/MALDI-TOF to assess the different forms of M^pro^. The eluting protease was analyzed by protein deconvolution (8.3-10 min) to determine the M^pro^ species present. For glutathionylated preparations made at pH 6.8 the M^pro^ was incubated for 15 min at 37°C in 50 mM Tris, pH 7.5, 2 mM EDTA, and 100 mM NaCl containing 100 ug/ml of cm-BSA, Grx (88-350 nM), 0.1 mM GSH or 0.1 mM GSH with 88-350 nM Grx in a total volume of 10 µL. After incubation an aliquot of each sample was assayed for M^pro^ activity (1 µM) and analyzed (2 µL) by SEC/MALDI-TOF to determine the percent of glutathionylation in each treatment based on the abundances of each species. For these experiments, the enzyme activity was assessed after stopping the reactions by acidification with FA/TFA and determining the pNA product produced using RP-HPLC, as described above, to quantitate the amount of pNA product generated over the 5 min incubation. TCEP treated glutathionylated enzyme was used to obtain the maximum native M^pro^ activity.

### Chymotrypsin and trypsin/lysC digestion and analysis of native and glutathionylated M^pro^

Native M^pro^ and M^pro^ which was monoglutathionylated (∼30%) as described above was digested with chymotrypsin or trypsin/lysC using the Accumap™ low pH sample preparation with urea under nonreducing conditions (Promega). The free cysteines in the M^pro^ preparations (100 µg) were first alkylated with N-ethylmaleimide in 8 M urea for 30 min at 37°C. Complete alkylation of all cysteines of the native M^pro^ with NEM was verified by RP-HPLC/MS-TOF analysis. For chymotrypsin digestion the alkylated proteins were diluted to 1 M urea with 100 mM Tris and 10 mM CaCl2 buffer pH 8.0 (50 µg of protease in 57 µl added to 456 µl of buffer) and treated with 2.5 µg of chymotrypsin made fresh in 1 mM HCl. Samples were incubated overnight (18 hours) at 37°C before stopping the reactions with a final of 2% TFA to reach a pH of <3.0. For typsin/r-LysC digestions the alkylated proteins were digested with low pH resistant r-Lys-C for 1 hours at 37°C followed by continued digestion with AccuMAP™ Modified Trypsin and AccuMAP™ Low pH Resistant rLys-C for 3 hours, as described in the AccuMAP™ protocol. The peptide digests were then cleaned up using peptide desalting columns (ThermoFisher) following the manufacturer’s instructions. The desalted clarified peptide mixtures were then dried in a Thermo speed vacuum system and resuspended in RP-HPLC solvent A (water with 0.1% FA/0.02%TFA). Aliquots of the peptide digests were then analyzed without or with TCEP-Cl treatment (50 mM) to remove glutathione modifications and then were separated on a Vydac C18 column. For peptide analysis the starting conditions were 100% solvent A (water with 0.1% FA/0.02%TFA). Elution of peptides was done with a 1%/min solvent B (acetonitrile with 0.1% FA/0.02%TFA) gradient over the first 20 minutes followed by a 2%/min gradient over the next 10 minutes. The elution of peptides was monitored using UV absorbance at 205, 254, and 276 nm as well as MALDI-TOF detection. Peptide digests were analyzed without and with TCEP (for native M^pro^ see Figure S2A and Figure S2B for UV and TIC chromatograms respectively and for monoglutathionylated M^pro^ digests without see Figure S2C and Figure S2D for UV and TIC chromatograms respectively or with TCEP analysis see Figure S2E and Figure S2F for UV and TIC chromatograms respectively). Chymotrypsin digestion of alkylated M^pro^ is predicted to produce 10 alkylated cysteine-containing peptides in addition to 12 other non-cysteine containing peptides of 3 amino acids or more. The predicted monoisotopic molecular masses for these peptides and their glutathionylated forms were used to extract specific peptide ions from the TIC chromatograms and the masses found were further confirmed by monoisotopic deconvolution. When glutathionylated masses were found, we then searched for their native counterparts following TCEP reduction. We could locate 6 of the 10 predicted alkylated cysteine containing peptides (covering 7 of the 12 cysteines) following chymotrypsin digestion of M^pro^ (see Table S1 for a list of peptides found in supplemental material). In addition to the predicted cysteine containing peptides, based on chymotrypsin digestion, the masses for two other cysteine containing peptides were identified including a 151:159 peptide fragment (containing cys156) and a 305:306 peptide fragment (containing cys300). These were produced, presumably, as a result of incomplete digestion by chymotrypsin at the 154:155 and 305:306 predicted cleavage sites (see Table S1, 7b and 10b, respectively, in supplemental material). We also found molecular masses consistent with 10 other non-cysteine containing peptides generated by chymotrypsin digestion (see Table S1 in supplemental material).

Trypsin/lysC digests were analyzed by RP-HPLC/MALDI-TOF for both native (see Figure S4A for TIC chromatogram and S4B for UV chromatogram in supplemental material) and monoglutathionylated preparations before (see Figure S4C for TIC chromatogram and S4D for UV chromatogram in supplemental material) and after TCEP treatment (see Figure S4E for TIC chromatogram and S4F for UV chromatogram in supplemental material). Trypsin/lysC digestion is predicted to yield 7 cysteine-containing peptides and 5 of the 7 cysteine alkylated peptides were found by molecular mass extraction from the TIC obtained by RP-HPLC/MALDI-TOF (see Table S2 in supplemental material). In addition to the predicted cysteine containing peptides, the masses for two other cysteine containing peptides were identified including a 41:61 peptide, resulting from incomplete cleavage at the 60:61 trypsin cleavage site, and a mass consistent with the tryptic peptide 299:306 having undergone spontaneous formation of the pyroglutamate form of the peptide (see Table S2 in supplemental material). This is commonly seen among peptides with N-terminal glutamates^21^ and its retention time and mass were confirmed using a synthetic peptide standard that contained both native and pyroglutamate forms.

### RP-HPLC MS-TOF analysis

Samples from the colorimetric enzyme assay, as described above, were analyzed by RP-HPLC with an Agilent 1200 series chromatograph on a Vydac C18-column (218TP5205, Hesperia, CA). Samples were injected (25-45 µL) and pNA substrate, pNA product and native and modified forms of M^pro^ were eluted with a 2%/min acetonitrile gradient beginning with 95% solvent A (0.1% FA)/0.02% TFA) in HPLC/MS grade water and 5% solvent B (0.1% FA/0.02% TFA in acetonitrile). The 2% gradient continued for 30 minutes and then was ramped to 95% acetonitrile in 2 minutes followed by a 5-minute re-equilibration to the starting conditions. Elution of samples was monitored at 205 nm, 276 nm, 320 nm (for pNA substrate) and 390 nm (for pNA product) with an Agilent diode array detector followed by MS analysis with an Agilent 6230 time of flight MS configured with Jetstream. M^pro^ and its glutathionylated forms eluted between 24-26 minutes (approximately 57% acetonitrile). The mass of the protein was determined by protein deconvolution using Agilent’s Mass Hunter software. The TOF settings were the following: Gas Temperature 350°C, drying gas 13 L/min, nebulizer 55 psi, sheath gas temperature 350°C, fragmentor 145 V, and skimmer 65 V. The mass determination for peptides was done by deconvolution (resolved isotope) using Agilent Mass Hunter software (Agilent).

### Analysis of M^pro^ by SEC coupled with MALDI-TOF MS detection

Size exclusion chromatography (SEC) on native and glutathionylated forms of M^pro^ was carried out using BioSep SEC3000 column and subsequently a BioSep SEC2000 column (300 mm × 4.6 mm; Phenomenex, Torrance, CA, U.S.A.) with 25 mM ammonium formate buffer (pH 8.0) running buffer on a 1200 series HPLC–MS system (Agilent, Santa Clara, CA, U.S.A.). The isocratic flow rate was 0.35 ml · min^-1^ and M^pro^ samples were injected at 2 µl. Where indicated, cm-BSA was used as a carrier to help prevent nonspecific binding of protein during the analysis. Proteins eluting from the column were monitored using an Agilent 1100 series fluorescent detector connected in series with the Agilent 6230 MS-TOF detector. At high concentrations, M^pro^ eluted as a single peak with a tailing edge while at lower concentrations M^pro^ eluted as two peaks consistent with it behaving as a monomer dimer system. For the SEC3000 column the M^pro^ peaks eluted between 8.5-10 minutes while for the SEC2000 column peaks eluted between 7-8.5 minutes. The percent of different forms of M^pro^ was estimated by using the abundances of each species which can only provide an estimate due to variations in ionization potential for each M^pro^ species.

### Analytical ultracentrifugation

For analytical ultracentrifugation (AUC) a Beckman Optima XL-I analytical ultracentrifuge, with absorption optics, an An-60 Ti rotor and standard double-sector centerpiece cells was used. Sedimentation equilibrium measurements of authentic native M^pro^ and glutathionylated M^pro^ were used to determine the average molecular weight and dissociation constant (*K*_d_) for dimerization. M^pro^ was diluted into 50 mM Tris pH 7.5 buffer containing 2 mM EDTA and 300 mM NaCl buffer to 1 µM (6 ml total solution) and then was untreated or glutathionylated with 10 mM GSSG for 45 minutes in the same buffer. Both preparations were washed by passing through a 10 kDa cut-off Amicon membrane and washing 4 times with 50 mM tris buffer with 2 mM EDTA and 100 mM NaCl. The preparations were analyzed by RP-HPLC/MS and control contained native M^pro^ while the glutathionylated preparation had predominantly diglutathionylated protease (63%), as well as triglutathionylated protease (22%) and monoglutathionylated protease (15%) based on their relative abundances. There was no detectable native M^pro^ remaining in this glutathionylated preparation. Proteins were concentrated to 0.63 mg/ml in 50 mM tris buffer pH 7.5 with 2 mM EDTA and 100 mM NaCl. Samples (100 µl) were centrifuged at 20°C at 21,000 rpm (16h) and 45,000 (3h) overspeed for baseline. Data (the average of 8 – 10 scans collected using a radial step size of 0.001 cm) were analyzed using the standard Optima XL-I data analysis software v6.03.

### Statistical analysis

Statistical analyses were performed using two-tailed Student’s *t-test* (paired) on experiments with at least 3 biological replicates or using a two-way ANOVA followed by Šídak’s multiple comparison post hoc test. P-values less or equal to 0.05 were considered statistically significant, *<0.05, **<0.01 and ***<0.005.

## Acknowledgements

This research was supported in part by the Intramural Research Program of the National Institutes of Health, National Cancer Institute and National Institute of Arthritis and Musculoskeletal and Skin Diseases. We thank Rodney Levine (National Heart, Lung and Blood Institute) and John Mieyal (Case Western Reserve University) for helpful discussions during this work.

## Author Contributions

Conceptualization, D.A.D. and R.Y.; investigation, D. A. D., H. B., P.S., A.Y., H. J., P.T.W., S.H. and H.M; original draft writing, D.A.D. and R.Y.; writing-review and editing, all authors; Funding acquisition, H.M. and R.Y.

## Materials & Correspondence

address to David A. Davis and/or Robert Yarchoan

## Data availability

All data are available in the main text or the supplementary materials.

## Figure Legends

**Figure 1: Exposure of low concentrations of SARS-CoV-2 M**^**pro**^ **to oxidized glutathione results in glutathionylation and inhibition of activity**. (A,B) Activity of M^pro^ following a 30-minute pre-incubation of (A) 1.2 µM M^pro^ or (B) 18 µM M^pro^ pretreated with 2 mM or 10 mM oxidized or reduced glutathione. After preincubation, M^pro^ was assayed for protease activity at an equal final enzyme concentration (1 µM). (C,D) M^pro^ molecular masses found by protein deconvolution for M^pro^ eluting off of the C18 reverse phase column following the different treatments at (C) 1.2 µM and (D) 18 µM. The theoretical molecular mass of M^pro^ is 33796.48 and the deconvoluted molecular mass for controls in (C) and (D) was 33797.09 and 33,797.34, respectively, as determined using Agilent’s Mass Hunter software. The experimental masses are shown above each peak obtained by deconvolution. The native M^pro^ as well as the increases in masses indicative of glutathionylation are indicated for the addition of 1 (+Δ1), 2 (+Δ2), and 3 (+Δ3) glutathione moieties in the deconvolution profiles of GSSG-treated M^pro^. Observed increases were 304, 609, and 913 as compared to the predicted increases of 305.1, 610.2 and 915.3 for addition of 1, 2 or 3 glutathione’s, respectively. Based on the abundances, the estimated percent of monoglutathionylation in (C) at 2 mM GSSG was 45% and for 10 mM GSSG there was an estimated 11% mono, 50% di, and 35% tri-glutathionylation, respectively. In (D) after treatment with 2 mM GSSG there was <5% monoglutathionylation and after 10 mM GSSG there was an estimated 34% monoglutathionylation. For (A) and (B) the values shown are the mean and standard deviation for three independent experiments (n=3) while for (C) and (D) the analysis was one time. (*** = p-value < 0.005, paired Students *t-test*). All other comparisons to control activity were not found to be significant p-value >0.05). M^pro^ control activity for (A) was 6.42 +/− 2.5 µM/min/mg and for (B) was 9.6 µM/min/mg, and the percent activity in the treatments is normalized to their respective controls.

**Figure 2: Inhibition of M**^**pro**^ **by glutathionylation can be reversed with reducing agent**. M^pro^ was glutathionylated at pH 7.5 with 10 mM GSSG as described in the Materials and Methods and the extent of glutathionylation was determined by RP/HPLC/MALDI-TOF using a 2% acetonitrile gradient as described in materials and Methods. (A,B) Deconvoluted masses obtained by protein deconvolution of the M^pro^ peak (eluting between 24 and 26 min) for (A) 3 µg (5 µL injection) purified glutathionylated M^pro^ and (B)) 3 µg (5 µL injection) glutathionylated M^pro^ after a 30 min treatment with 10 mM DTT. Shown above each peak is the molecular mass (top number) and the abundance (bottom number) found by protein deconvolution. The native, monoglutathionylated (+Δ1), diglutathionylated (+Δ2), and triglutathionylated (+Δ3) M^pro^, are indicated in the figures. (C) M^pro^ activity (1 µM final enzyme) for native and glutathionylated M^pro^ preparations after a 30-min incubation in the absence or presence of 10 mM DTT. M^pro^ activity for control in (C) and was 4.95 +/− 1.2 µM/min/mg and percent activity for the different conditions was normalized to their respective controls. The values shown are the average and standard deviation from three separate experiments (n=3) (* = p-value < 0.05, paired students *t-test*, ns = not significant).

**Figure 3: Size exclusion chromatography and equilibrium analytical ultracentrifugation of M**^**pro**^ **and glutathionylated M**^**pro**^ **indicates glutathionylated M**^**pro**^ **behaves as a monomer**. (A,D) M^pro^ and glutathionylated M^pro^ were analyzed by SEC3000/MALDI-TOF. (A) Overlay of the chromatograms for 60 µM each of M^pro^ (black line) and glutathionylated M^pro^ (red line) and (D) 7.5 µM each of M^pro^ (black line) and glutathionylated M^pro^ (red line) by monitoring the intrinsic protein fluorescence (excitation 276 nm, emission 350 nm). Glutathionylated M^pro^ was made with 10 mM GSSG at pH 7.5 for 2-2.5 hours as described in Materials and Methods. (C,D) Protein deconvolution profiles for (B) native M^prov^ and (C) glutathionylated M^pro^ that were run as shown in (A). (E,F) Protein deconvolution profile for (E) native M^pro^ and (F) glutathionylated M^pro^ that were run as shown in (D). Shown above each peak are the molecular mass (top number) and the abundance (bottom number) found by protein deconvolution. The earlier eluting peak at 8.5 min is cm-BSA, which was used as a carrier in the runs of M^pro^ to help prevent potential non-specific losses of protein during the run. (G,H) Equilibrium analytical ultracentrifugation of (G) M^pro^ and (H) glutathionylated M^pro^ at 0.63 mg ml^-1^ (18 µM) in 50 mM tris buffer pH 7.5, 2 mM EDTA, and 100 mM NaCl. The absorbance gradients in the centrifuge cell after the sedimentation equilibrium was attained at 21,000 rpm are shown in the lower panels. The open circles represent the experimental values, and the solid lines represent the results of fitting to a single ideal species. The best fit for the data shown in (G) yielded a relative molecular weight (*Mr*) of 62,991 +/− 1144 and a K_d_ for dimerization of 2.4 µM and that shown in (H) yielded a molecular weight of 37,000 +/− 1000 and a K_d_ for dimerization of 200 µM. The corresponding upper panels show the differences in the fitted and experimental values as a function of radial position (residuals). The residuals of these fits were random, indicating that the single species model is appropriate for the analyses.

**Figure 4: Size exclusion chromatography of a preparation of monoglutathionylated M**^**pro**^ **and analysis of M**^**pro**^ **activity**. A preparation of M^pro^ containing a mixture of native and monoglutathionylated forms was made by incubating 1.2 µM M^pro^ with 5 mM GSSG for 2.5 hours at 37°C, at pH 6.8, to increase specific modification of the more reactive cysteines of M^pro^ as described in Materials and Methods. (A) SEC2000 elution profile as monitored using the intrinsic protein fluorescence (excitation 276 nm, emission 350 nm) of a 2 µl injection of 8 µM monoglutathionylated M^pro^ preparation and (B) M^pro^ molecular weights found by protein deconvolution of the peaks in (A), (C) Elution profile for the mass of native M^pro^ in the monoglutathionylated preparation and (D) elution profile for the mass of monoglutathionylated M^pro^ in the monoglutathionylated preparation. (E) Elution profile for 2 µl injection of 8 µM monoglutathionylated M^pro^ preparation after treatment with 50 mM TCEP for 15 min. (F) M^pro^ molecular weights found by protein deconvolution after treatment with 50 mM TCEP for 15 min. (G) Elution profile for the mass of native M^pro^ after treatment of monoglutathionylated M^pro^ with 50 mM TCEP for 15 min. (H) M^pro^ activity without (black bars) and with (grey bars) TCEP treatment for peak #1 and Peak #2 from Fig 4A after collecting M^pro^ following SEC of the monoglutathionylated M^pro^ preparation. The values represent the average of 4 separate determinations (n=4) of M^pro^ activity. A two-way ANOVA followed by Šídak’s multiple comparison post hoc test was done. P-values less or equal to 0.05 were considered statistically significant, **<0.01 and ***<0.005 (ns= p-value > 0.05).

**Figure 5: Grx reverses glutathionylation and restores M**^**pro**^ **activity**. (A-C) M^pro^ Glutathionylated at pH 7.5 was incubated (3 µM final) for 30 min in the presence of (A) buffer control, (B) GSH (0.5 mM) or (C) GSH (0.5 mM) with Grx (final 350 nM). Samples were analyzed by SEC3000/MALDI-TOF and the eluting protease analyzed by protein deconvolution (8.3-10 min) to determine the M^pro^ species present. The experimental masses (top number) are shown as well as the abundances (bottom number) for each peak obtained by deconvolution. The native M^pro^, as well as the increases in masses indicative of glutathionylation, are indicated for the addition of 1 (+Δ1), 2 (+Δ2), and 3 (+Δ3) glutathione moieties in the deconvolution profiles. (D) Samples of glutathionylated M^pro^ were treated as in (A-C) and then analyzed for M^pro^ activity and compared to unmodified M^pro^. M^pro^ activity for control in (D) was 5.77+/− 1.5 µM/min/mg, and percent activity for the different conditions was normalized to their respective controls. (E-G) Monoglutathionylated M^pro^ was incubated (8 µM final) for 15 min in the presence of (A) buffer control, (B) GSH (0.1 mM) or (C) GSH (0.1 mM) with Grx 350 nM and samples analyzed by SEC2000/MALDI-TOF deconvolution (7.3-8.6 min). (H, I) Samples were prepared as in (E-G) and the percentage of monoglutathionylated M^pro^ and activity was determined after the 15-minute incubation with 0, 88, 175, or 350 nm Grx in the presence of 100 µM GSH. (H) Percent of monoglutathionylated Mpro after Grx treatment and (I) M^pro^ activity after Grx treatment. The M^pro^ activity was normalized to the TCEP treated preparation which yielded fully reduced native M^pro^ and was used as 100% activity. For (D) Values represent the average +/− standard deviation of 4 separate experiments (* = p-value < 0.05, ****=p-value < 0.001 paired students t-test, ns=not significant p>0.05). For (H) the values are the average of 3 separate experiments (n=3) and for (I) one experiment performed in duplicate (n=2).

**Figure 6: Glutathionylation inhibits WT SARS-Cov-2 M**^**pro**^ **activity but not C300S M**^**pro**^ **activity**. (A) Activity of wild type (WT) and C300S M^pro^ (1 µM enzyme) following a 30-minute pre-incubation of 1.2 µM M^pro^ with 10 mM oxidized glutathione. (B) M^pro^ activity for a WT monoglutathionylated M^pro^ preparation (containing approximately 30% monoglutathionylated M^pro^ and 4% diglutathionylated) and a C300S monoglutathionylated M^pro^ preparation (containing approximately 18% monoglutathionylated M^pro^) preincubated for 10 minutes without or with 20 mM DTT. The amount of monoglutathionylated M^pro^ was estimated using the relative abundances of native M^pro^ and glutathionylated M^pro^ following deconvolution of the eluting M^pro^ species from SEC/MALDI-TOF analysis. Values represent the average +/− standard deviation of 3 separate experiments (n=3) (* = p-value < 0.05, ***=p-value < 0.005 paired students t-test, ns=not significant p>0.05).

**Figure 7: The current model for the regulation of dimerization and activity through reversible glutathionylation of M**^**pro**^ **and Space filling and close up ribbon model of SARS-CoV-2 M**^**pro**^. (A) Model showing that M^pro^ dimer exists in equilibrium with its monomer form with a determine K_d_ of 2.5 µM. The monomeric M^pro^ is susceptible to glutathionylation at Cys300, and this leads to inhibition of dimerization and loss of activity. Human Grx is able to reverse glutathionylation of Cys300 and restore dimerization and activity. (B) Space filling model of the SARS-CoV-2 M^pro^ dimer (apo form) showing the location of cysteines 156 on the surface and 300 near the dimer interface in the left (pink) protomer (PDB ID 7K3T). (C) Close up ribbon model around Cys300 showing the proximity to protomer 2 (blue) at leucine 141’ and the proximity to ASN214, GLN299 and PHE3.

**Figure 8: The local environment around Cys300 in monomeric SARS-CoV-1 M**^**pro**^. Ball and stick model for local environment around cys300 in R298A M^pro^ monomer PDB ID 2QCY (a monomeric form of SARS-CoV M^pro^ mutant R298A at pH 6.0). Structural figures were produced with PyMOL v1.5.0.4 ^40^.

