## Supplemental Figs and Tables for "Regulation of the Dimerization and Activity of SARS-CoV-2 Main Protease through Reversible Glutathionylation of Cysteine 300"

#### Supplementary data

**Figure S1: Analysis of purified WT and C300S $M^{pro}$ by SDS-gel electrophoresis and RP-HPLC/MALDI-TOF analysis.** (A) Amino acid sequence for WT $M^{pro}$. The arrow indicates cysteine 300 which was mutated to serine300 in the mutant Mpro. Shown below the sequence is the calculated molecular weight ($M_r$) for WT and C300S Mpro. (B,F) LDS-gel electrophoresis of (B) purified WT $M^{pro}$ and (F) purified C300S $M^{pro}$. In each gel, lanes 1 and 4 are molecular weight markers while lanes 2 and 5 are 4 µg and 8 µg of total protein loaded, respectively. (C,G) RP/HPLC showing the UV chromatogram at 205nm for (B) WT and (G) C300S $M^{pro}$. (D,H) The MALDI-TOF TIC chromatogram obtained by mass spectrometry for (D) WT and (H) C300S $M^{pro}$. (E,I) Protein deconvolution of the peaks in (D) and (H) showing the determined molecular weight obtained for (E) purified $M^{pro}$ (33797.5 experimental vs 33796.5 calculated) and (I) C300S $M^{pro}$ (33781.8 experimental vs 33780.4 calculated) after protein deconvolution. The insets in (E) and (I) show the molecular ion profile used for deconvolution of each $M^{pro}$. Separations were done on a Vydac C18 column using 95% buffer A (water 0.1% formic acid and 0.02% trifluoroacetic acid) and 5% buffer B (acetonitrile with 0.1% formic acid and 0.02% trifluoroacetic acid) with a 0.5 ml $min^{-1}$ flow rate and ramped to 65% buffer B with a 2% gradient for 30 minutes and then ramped to 100% for the next 5 min and then returned to starting conditions 2 min later. $M^{pro}$ eluted at approximately 25 min. The TOF settings were as follows: gas temperature, 350°C; drying gas rate, 13 liters/min; nebulizer, 55 pounds per square inch gauge (psig); sheath gas temperature, 350°C; fragmenter, 350 V; skimmer, 65 V. Molecular weights were determined by protein deconvolution using Agilent Mass Hunter software (Agilent).

**Figure S2: RP-HPLC/MALDI-TOF and UV chromatograms of chymotryptic digestions of native and monoglutathinylated $M^{pro}$ preparations.** Native and monoglutathionylated $M^{pro}$ preparations were alkylated and digested with chymotrypsin as described in the materials and methods and then anlayzed by RP-HPLC/MALDI-TOF for the identification of $M^{pro}$ peptides. (A) TIC chromatogram and (B) UV chromatogram (205 nm) of the native $M^{pro}$ peptide digest. (C) TIC chromatogram and (D) UV chromatogram (205 nm) of the monoglutathionyated $M^{pro}$ peptide digest. (E) TIC chromatogram and (F) UV chromatogram (205 nm) of the monoglutathionyated $M^{pro}$ peptide digest following treatment with TCEP to reduce disulfide bonds.

**Figure S3: Identification of glutathionylation at cysteines 156 and 300 in chymotryptic digests of monoglutathionylated $M^{pro}$ preparations.** The monoglutathionylated $M^{pro}$ preparation was alkylated and digested with chymotrypsin as described in the materials and methods and then analyzed by RP-HPLC/MALDI-TOF for the identification of $M^{pro}$ peptides. (A,B) The relevant area of the RP-HPLC/MS TIC chromatogram for the elution of chymotryptic peptides before (A) or after (B) treatment of the digested peptide preparation with TCEP to remove glutathione from glutathionylated peptides. (C,D) The corresponding RP-HPLC/MS UV chromatogram at 205 nm for the elution of chymotryptic peptides before (C) or after (D) treatment of the digested peptide preparation with TCEP to remove glutathione from glutathionylated peptides. The arrows in (A) and (C) indicate the location of eluting glutathionylated peptides $1^g$,$2^g$ and $3^g$ and in (B) and (D) indicate the location for the corresponding native peptides $1^n$,$2^n$ and $3^n$ detected after reduction with TCEP. SSG denotes glutathionylated peptide. (E, G, I) Identification of peak $3^g$ (E), peak $2^g$ (G) and peak $1^g$ (I) by ion extraction followed by monoisotopic deconvolution revealing masses corresponding to the glutathionylated chymotryptic peptides; 151:159, 295:305 and 295:306. Each chromatogram shows the extracted ion chromatogram using the predicted monoisotopic masses $[M+H]^{+1}$ for 151:159-SSG, 295:305-SSG and 295:306-SSG, respectively. (E,G,I) The insets in E, G, and I show the deconvoluted monoisotopic masses obtained for the peak with the monoisotopic masses indicated by dashed arrows. (F, H, and J) the same analysis as in E,G and I showing the absence of the glutathionylated peptides after TCEP reduction. (L,N, and P) Identification of peak $3^n$ (L), peak $2^n$ (N) and peak $1^n$ (P) by ion extraction followed by monoisotopic deconvolution revealing native peptides; 151:159, 295:305 and 295:306. Each chromatogram shows the extracted ion chromatogram using the predicted monoisotopic masses $[M+H]^{+1}$ for 151:159, 295:305 and 295:306, respectively. The insets in (L, N, and P) show the deconvoluted monoisotopic masses obtained for each peptide with the monoisotopic masses indicated by dashed arrows. For (K, M and O) the same analysis is done on the TCEP treated sample which reveals the loss of detection of the glutathionylated peptides. (L, N, and P) and identification of the native peptides (L) 151:159 (N) 295:305 and (P) 295:306 following TCEP treatment. Each panel shows the extracted ion chromatogram for the predicted monoisotopic masses 151:159, 295:305 and 295:306, respectively. The insets in (L, N, and P) show the deconvoluted monoisotopic masses obtained for each peptide with the monoisotopic masses indicated by dashed arrows. For (K, M and O) the same native molecular peptide ion extraction analysis is done on the bnon TCEP treated sample to show the absence of these peptides in the sample prior to TCEP treatment.

**Figure S4: RP-HPLC/MALDI-TOF and UV chromatograms of trypsin/lysC digestions of native and monoglutathinylated $M^{pro}$ preparations.** Native and monoglutathionylated $M^{pro}$

preparations were alkylated and digested with trypsin/lysC as described in the materials and methods and then anlayzed by RP-HPLC/MALDI-TOF for the identification of $M^{pro}$ peptides. (A) TIC chromatogram and (B) UV chromatogram (205 nm) of the native $M^{pro}$ peptide digest. (C) TIC chromatogram and (D) UV chromatogram (205 nm) of the monoglutathionyated $M^{pro}$ peptide digest. (E) TIC chromatogram and (F) UV chromatogram (205 nm) of the monoglutathionyated $M^{pro}$ peptide digest following treatment with TCEP to reduce disulfide bonds.

**Figure S5: Identification of Cys300 as a major target for glutathionylation based on trypsin/lysC digests of Monoglutathionylated $M^{pro}$ preparations.**The monoglutathionylated $M^{pro}$ preparation was alkylated and digested with trypsin/lysC as described in the materials and methods and then analyzed by RP-HPLC/MALDI-TOF for the identification of $M^{pro}$ peptides. (A,B) The relevant area of the RP-HPLC/MS TIC chromatogram for the elution of trypsin/lysC generated peptides before (A) or after (B) treatment of the digested peptide preparation with TCEP to remove glutathione from glutathionylated peptides. (C,D) RP-HPLC/MS UV chromatogram at 205 nm for the elution of trypsin/lysC generated peptides before (C) or after (D) treatment of the digested peptide preparation with TCEP to remove glutathione from glutathionylated peptides. The arrows in A and C indicate the location of eluting glutathionylated peptides 1,2 and 3 and in B and D indicate the location for the native peptides 1`,2` and 3` detected after reduction with TCEP. SSG denotes glutathionylated peptide and (py) denotes the pyroglutamate form of the 299-306 peptide that results from spontaneous deamidation of peptides with N-terminal glutamyl residues {Wright, 1991 #28}. (E-I) Detection and identification of glutathionylated typ/lysC peptides 77:88-SSG, (G) 299:306-SSG py (pyroglutamte form of the peptide) and (I) 295:306-SSG peptides. Each panel in E, G, and I shows the extracted ion chromatogram for the predicted monoisotopic masses for 77:88-SSG, 299:306-SSGpy, and 299:306-SSG, respectively. The insets in E, G, and I show the deconvoluted monoisotopic masses obtained for each glutathionylated peptide with the monoisotopic masses indicated by dashed arrows. (F-J) the same analysis is done on the TCEP treated sample which reveals the loss of detection of the glutathionylated peptides when carrying out the same mass extractions as E, G, and I. (L, N, and P) Detection and Identification of the native peptides (L) 77:88 (N) 299:306py and (P) 295:306 following TCEP treatment. Each panel shows the extracted ion chromatogram for the predicted monoisotopic masses for peptides 77:88, 299:306py and 299:306, respectively. The insets in L, N, and P show the deconvoluted monoisotopic masses obtained for each peptide with the monoisotopic masses indicated by dashed arrows. For K, M and O the same same analysis is done on the untreated sample which shows the absence of the peaks seen in L,N and P prior to TCEP treatment.

**Figure S6: Glutathionylation of native and C300S M$^{pro}$ after treatment with GSSG.** Masses found by protein deconvolution from RPHPLC/MALDI -TOF analysis of samples in Figure 6A. (A) WT M$^{pro}$ or (B) WT M$^{pro}$ treated with 10 mM GSSG for 30 min. Masses found by protein deconvolution for (C) C300S M$^{pro}$ or (D) C300S M$^{pro}$ after treatment with 10 mM GSSG for 30 min. Samples were analyzed for protein glutathionylation following the assays performed in Figure 6A. (E) SEC/MALDI-TOF analysis of WT and (F) C300S M$^{pro}$ after glutathionylating at pH 6.8 with 5 mM GSSG as described in material and methods. The upper number above each peak denotes the calculated mass, and the lower number denotes the abundance.

**Supplemental Figure S7**. Comparison of the local environment around Cys300 in dimeric SARS-CoV-1 Mpro and dimeric SARS-CoV-2 Mpro. (A) Ball and stick model for local environment around cys300 in SARS-CoV-1 Mpro showing the interactions with ASN214 and ASN299. Mpro (PDB ID 1UJ1 (SARS-CoV-1 Mpro apoenzyme at pH 6.0). (B) Ball and stick model for local environment around cys300 in SARS-CoV-2 Mpro showing the interactions with ASN214 and ASN299 carbonyls (PDB ID 7KT (SARS-CoV-2 Mpro apoenzyme at pH 6.5). Structural figures were produced with PyMOL v1.5.0.4 40.

**Supplemental Figure S8**. Amino acid alignment of the Main Protease from bat isolates (document ID's are MN996532.2 for RaTG13, MG772933.1 for ZC45, MG772934.1 for ZXC21 .nd MW681072.1 for SARS-CoV-2). Arrows indicate amino acids that are not fully conserved among all 4 sequences. There are two amino acid differences between SARS CoV-2 and RatG13 and 3 amino acid differences between SARS CoV-2 and both CoV-ZC45 and CoV-ZXC and those differences are highlighted in grey. Cysteine 300 is shown in the box.

### Supplemental Figures for

#### Regulation of the Dimerization and Activity of SARS-CoV-2 Main Protease through Reversible Glutathionylation of Cysteine 300

by Davis et al.

#### Supplemental Figure 1

A

amino acid sequence of WT $M^{pro}$

```
SGFRKMAFPSGKVEGCMVQVTCGTTTLNGLWLDDVVYCPRHVICTSEDMLNPNYEDLLIRKSNHNFLVQAGNVQLRVIGHSMQN
CVLKLKVDTANPKTPKYKFVRIQPGQTFSVLACYNGSPSGVYQCAMRPNFTIKGSFLNGSCGSVGFNIDYDCVSFCYMHHMELP
TGVHAGTDLEGNFYGPFVDRQTAQAAGTDTTITVNVLAWLYAAVINGDRWFLNRFTTTLNDFNLVAMKYNYEPLTQDHVDILGP
LSAQTGIAVLDMCASLKELLQNGMNGRTILGSALLEDEFTPFDVVRQCSGVTFQ
```

WT $M^{pro}$ Calculated molecular weight average ($M_{r\ (calc)}$): 33796.48 amu

C300S $M^{pro}$ Calculated molecular weight average ($M_{r\ (calc)}$): 33780.42 amu

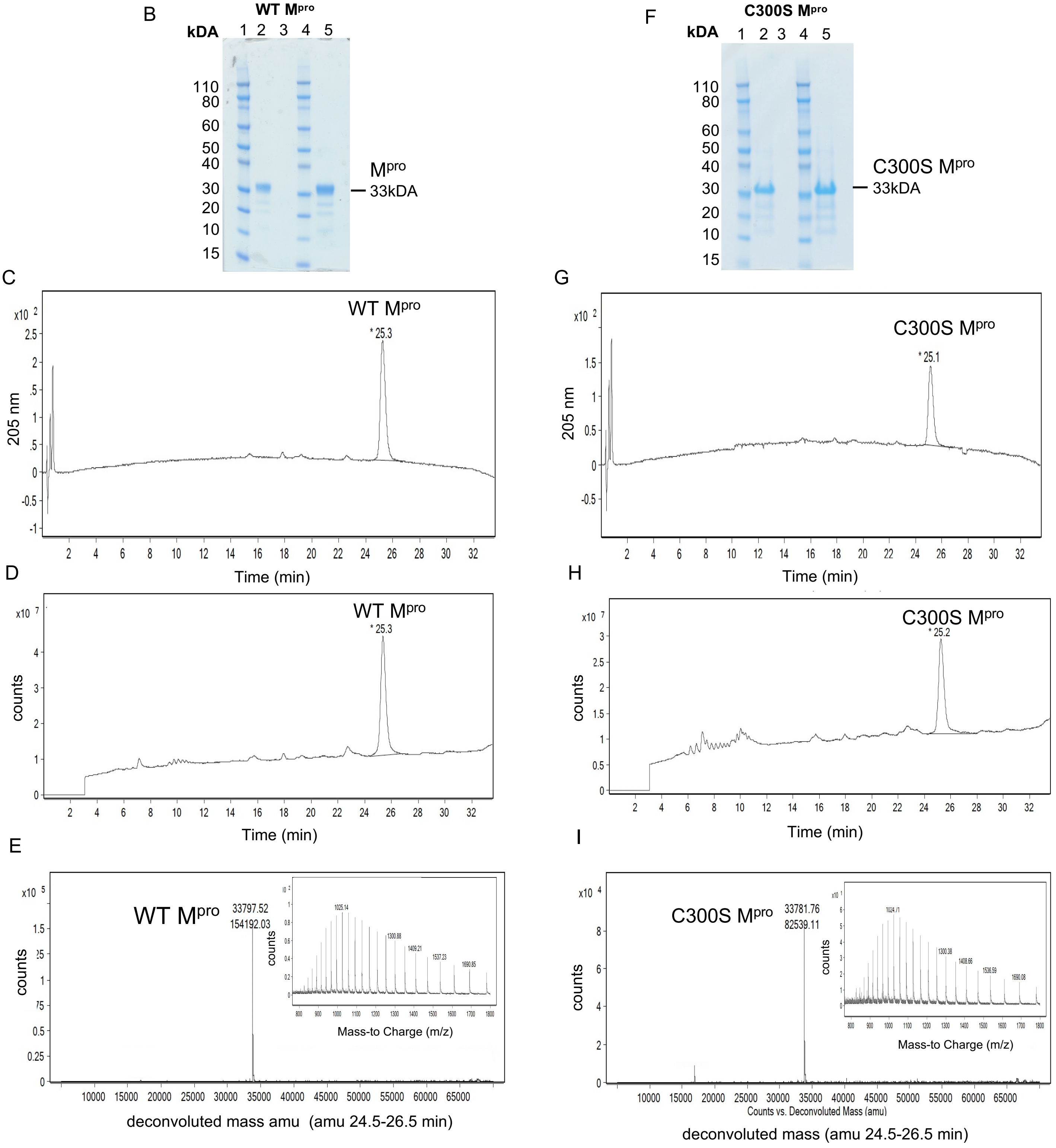

### Supplemental Figure 2

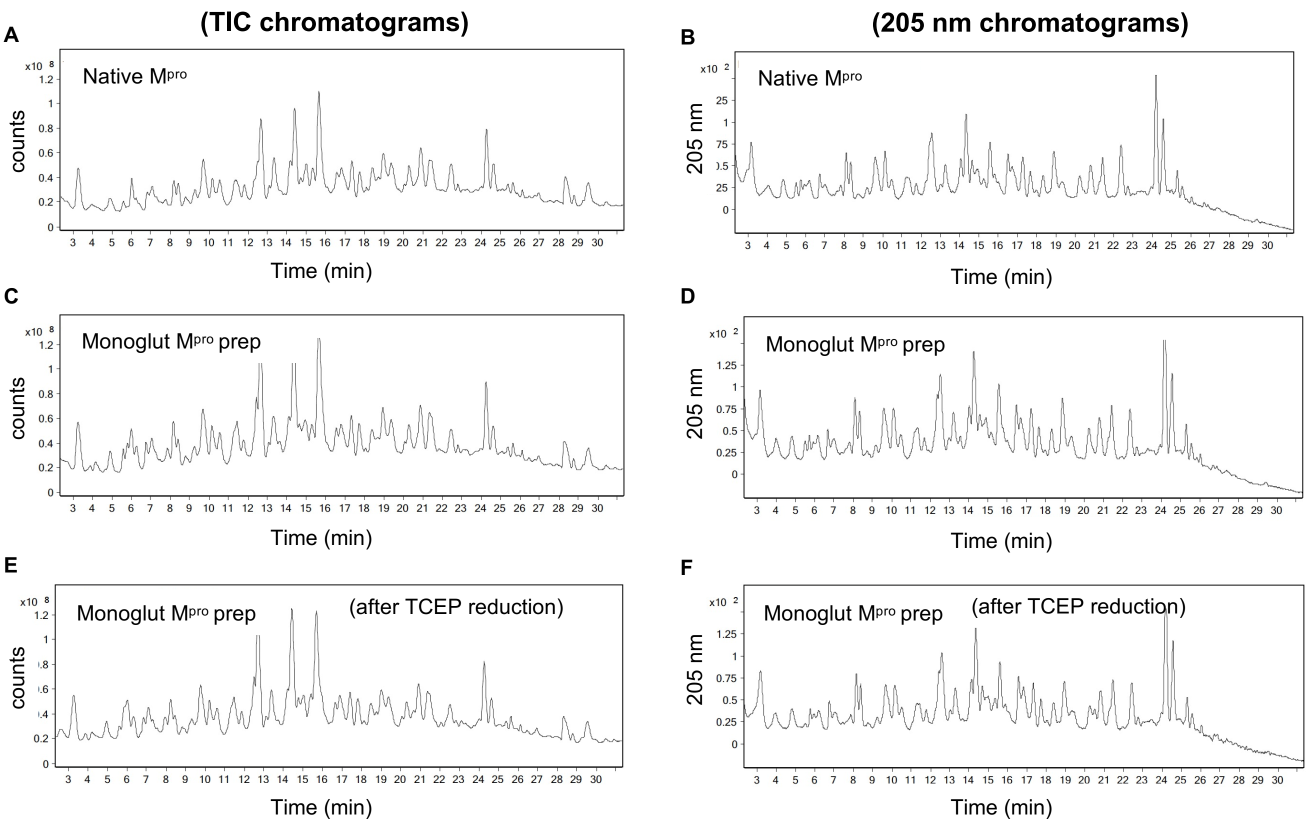

### Supplemental Figure 3

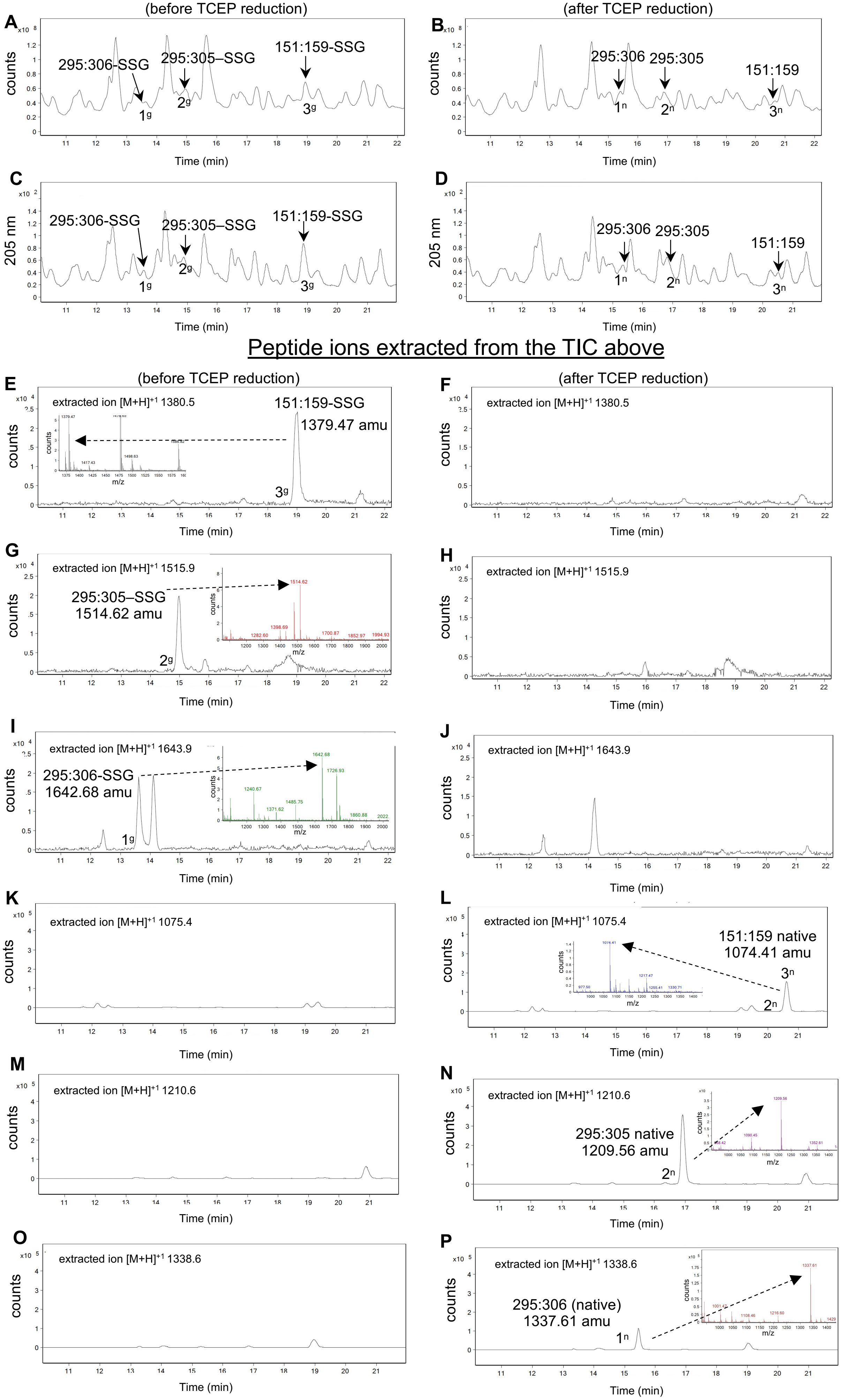

### Supplemental Figure 4

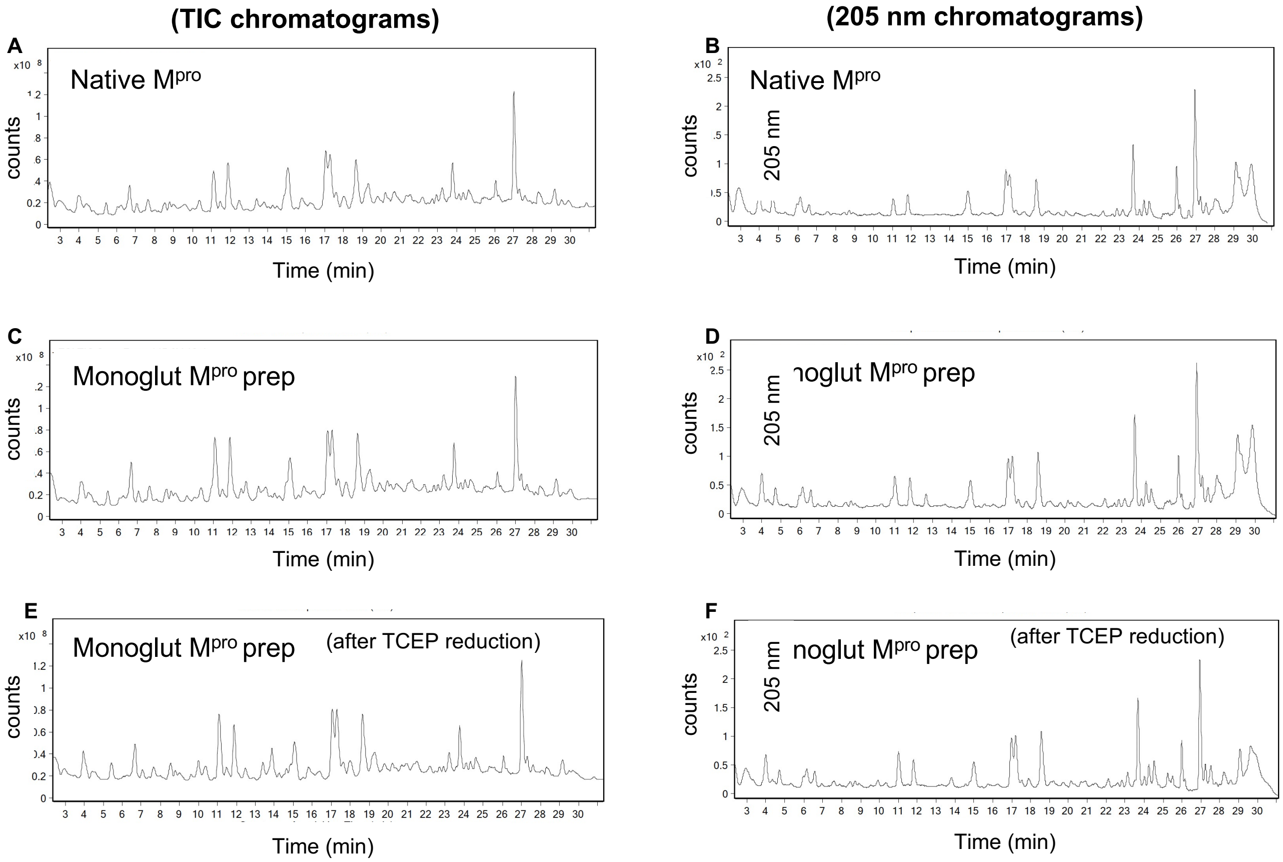

### Supplemental Figure 5

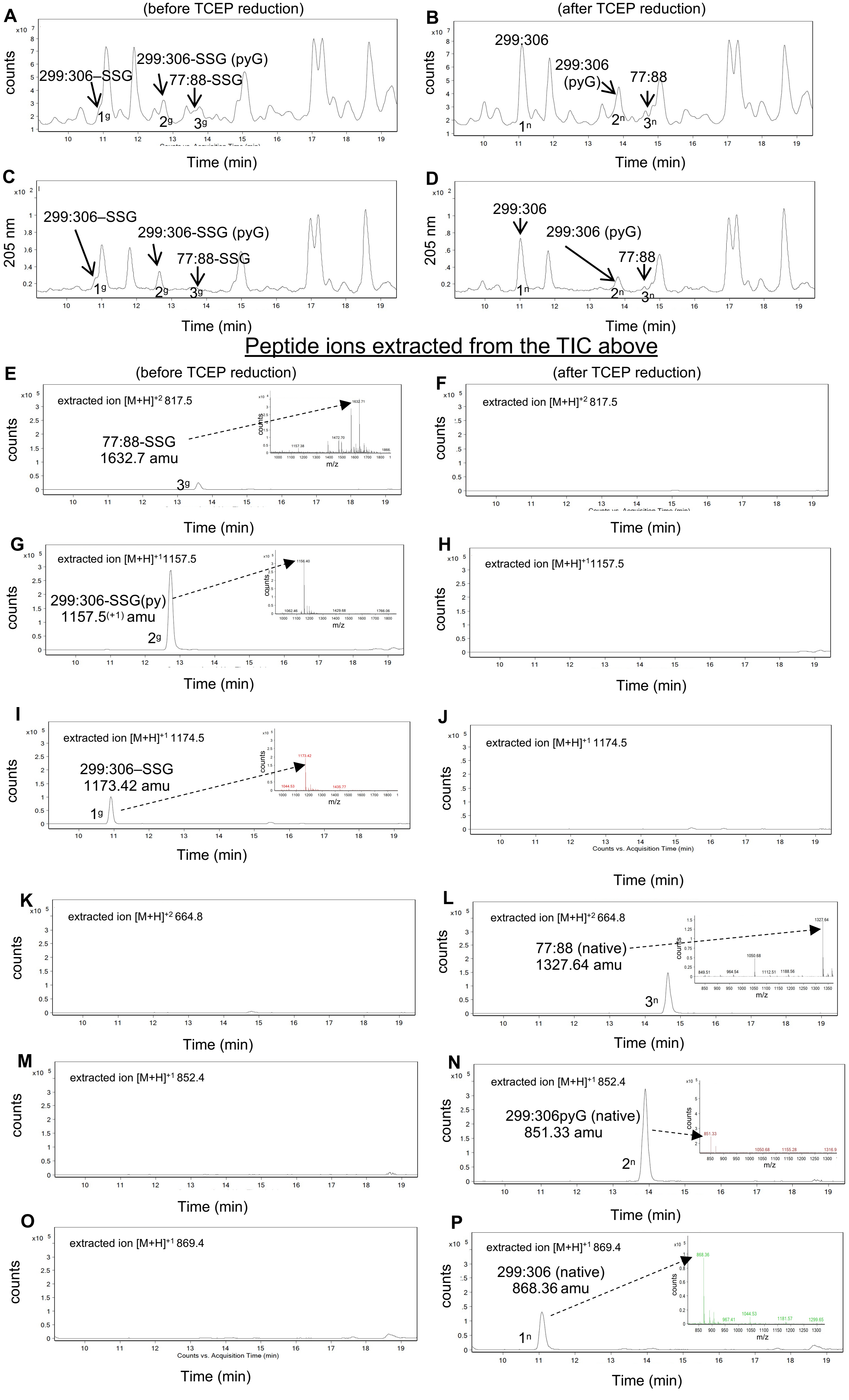

### Supplemental Figure 6

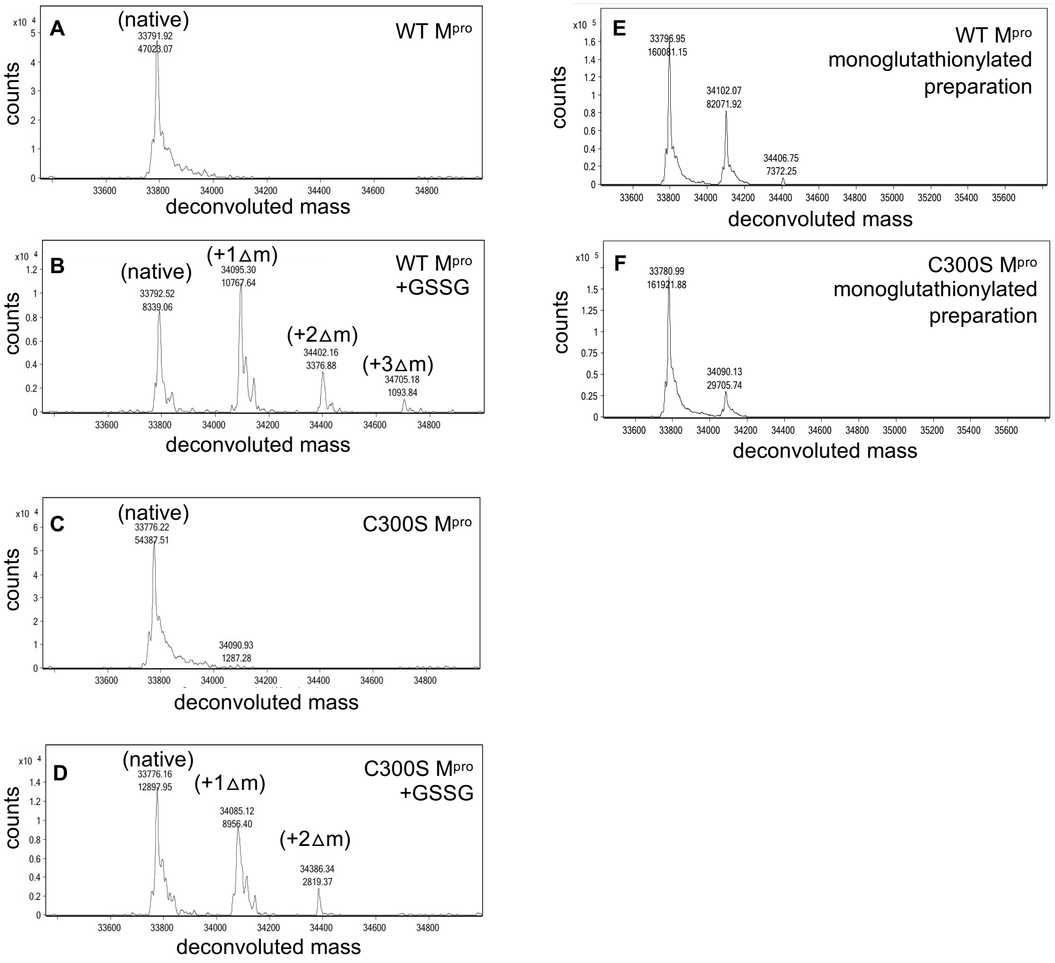

### Supplemental Figure 7

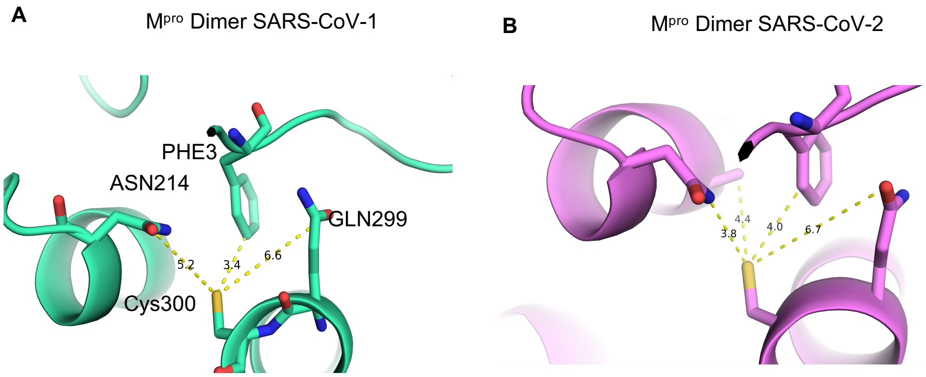

#### Supplemental Figure 8

```
Bat-SL-CoV-ZC45   1 SGFRKMAFPSGKVEGCMVQVTCGTTTLNGLWLDDVVYCPRHVICTSEDMLNPNYEDLLIR  60
Bat Mpro CoVZXC   1 SGFRKMAFPSGKVEGCMVQVTCGTTTLNGLWLDDVVYCPRHVICTSEDMLNPNYEDLLIR  60
RaTG13 Mpro QHR   1 SGFRKMAFPSGKVEGCMVQVTCGTTTLNGLWLDDVVYCPRHVICTSEDMLNPNYEDLLIR  60
SARS2 Mpro prot   1 SGFRKMAFPSGKVEGCMVQVTCGTTTLNGLWLDDVVYCPRHVICTSEDMLNPNYEDLLIR  60
                    ************************************************************

Bat-SL-CoV-ZC45  61 KSNHNFLVQAGNVQLRVVGHSMQNCVLKLKVDTANPKTPKYKFVRIQPGQTFSVLACYNG 120
Bat Mpro CoVZXC  61 KSNHNFLVQAGNVQLRVVGHSMQNCVLKLKVDTANPKTPKYKFVRIQPGQTFSVLACYNG 120
RaTG13 Mpro QHR  61 KSNHNFLVQAGNVQLRVIGHSMQNCVLKLKVDTANTKTPKYKFVRIQPGQTFSVLACYNG 120
SARS2 Mpro prot  61 KSNHNFLVQAGNVQLRVIGHSMQNCVLKLKVDTANPKTPKYKFVRIQPGQTFSVLACYNG 120
                    ****************.***************** *************************

Bat-SL-CoV-ZC45 121 SPSGVYQCAMRPNFTIKGSFLNGSCGSVGFNIDYDCVSFCYMHHMELPTGVHAGTDLEGT 180
Bat Mpro CoVZXC 121 SPSGVYQCAMRPNFTIKGSFLNGSCGSVGFNIDYDCVSFCYMHHMELPTGVHAGTDLEGT 180
RaTG13 Mpro QHR 121 SPSGVYQCAMRPNFTIKGSFLNGSCGSVGFNIDYDCVSFCYMHHMELPTGVHAGTDLEGT 180
SARS2 Mpro prot 121 SPSGVYQCAMRPNFTIKGSFLNGSCGSVGFNIDYDCVSFCYMHHMELPTGVHAGTDLEGN 180
                    ***********************************************************.

Bat-SL-CoV-ZC45 181 FYGPFVDRQTAQAAGTDTTITVNVLAWLYAAVINGDRWFLNRFTTTLNDFNLVAMKYNYE 240
Bat Mpro CoVZXC 181 FYGPFVDRQTAQAAGTDTTITVNVLAWLYAAVINGDRWFLNRFTTTLNDFNLVAMKYNYE 240
RaTG13 Mpro QHR 181 FYGPFVDRQTAQAAGTDTTITVNVLAWLYAAVINGDRWFLNRFTTTLNDFNLVAMKYNYE 240
SARS2 Mpro prot 181 FYGPFVDRQTAQAAGTDTTITVNVLAWLYAAVINGDRWFLNRFTTTLNDFNLVAMKYNYE 240
                    ************************************************************

Bat-SL-CoV-ZC45 241 PLTQDHLDILGPLSAQTGIAVLDMCASLKELLQNGMNGRTILGSALLEDEFTPFDVVRQC 300
Bat Mpro CoVZXC 241 PLTQDHLDILGPLSAQTGIAVLDMCASLKELLQNGMNGRTILGSALLEDEFTPFDVVRQC 300
RaTG13 Mpro QHR 241 PLTQDHVDILGPLSAQTGIAVLDMCASLKELLQNGMNGRTILGSALLEDEFTPFDVVRQC 300
SARS2 Mpro prot 241 PLTQDHVDILGPLSAQTGIAVLDMCASLKELLQNGMNGRTILGSALLEDEFTPFDVVRQC 300
                    ******.*****************************************************

Bat-SL-CoV-ZC45 301 SGVTFQ 306
Bat Mpro CoVZXC 301 SGVTFQ 306
RaTG13 Mpro QHR 301 SGVTFQ 306
SARS2 Mpro prot 301 SGVTFQ 306
                    ******
```

**Table S1: Peptides calculated and observed after cysteine alkylation and chymotrypsin digestion of $M^{pro}$ or monoglutathionylated $M^{pro}$ preparations**

| Peptide Number | Peptide From:To | $M_{r\ (calc)}$ | $M_{r\ (expt)}$ | Delta | R.T. |
|---|---|---|---|---|---|
| 1 | 9:31 (cys16 & 22) | 2630.22 | ND | - | - |
| 2 | 38:54 (cys38 & 44) | 2240.96 | 2239.41. | 1.55 | 24 |
| 3 | 67:101 (cys85) | 3987.16 | ND | - | - |
| 41 | 113:118 (cys117) | 779.35 | 779.33 | -0.02 | 17.7 |
| 5a[1] | 127:134 (cys128) | 1090.47 | 1090.45 | -0.02 | 16.9/17.3 |
| 5b[1] | py127:134 (cys128) | 1073.47 | 1073.44 | -0.03 | 18.9/19.3 |
| 61 | 141:150 (cys145) | 1064.46 | 1064.44 | -0.02 | 18.1 |
| 7a[1] | 155:159 (cys156) | 694.26 | 694.36 | 0.1 | 9.7 |
| 7b[1] | 151:159 (cys156)* | 1199.43 | 1199.46 | 0.03 | 22.5 |
| 8 | 160:161 (cys161) | 409.13 | ND | - | - |
| 9 | 240:291 (cys265) | 5560.84 | ND | - | - |
| 10a[1] | 295:305 (cys300) | 1334.63 | 1334.61 | -0.02 | 19 |
| 10b[1] | 295:306 (cys300)** | 1462.69 | 1462.66 | -0.03 | 17.4 |
| 111 | 141:150 (cys-sg 145) | 1244.48 | 1240.67 | -3.81 | 13.7 |
| 11n[1] | 141:150 (native) | 939.4 | 939.4 | 0 | 15.1 |
| 121 | 151:159 (cys-sg 156) * | 1379.3 | 1379.47 | 0.17 | 19 |
| 12n[1] | 151:159 (native) | 1074.42 | 1074.41 | -0.01 | 20.7 |
| 131 | 295:306 (cys-sg 300) | 1642.7 | 1642.68 | -0.02 | 13.7 |
| 13n[1] | 295:306 (native) ** | 1337.63 | 1337.61 | -0.02 | 15.5 |
| 141 | 295:305 (cys-sg 300) | 1514.66 | 1514.62 | -0.04 | 14.9 |
| 14n[1] | 295:305 (native) | 1209.58 | 1209.56 | -0.02 | 16.9 |
| 15 | 4:8 | 651.34 | 651.35 | 0.01 | 9.7*** |
| 16 | 32:37 | 722.34 | 722.34 | -0.02 | 13.3 |
| 17 | 55:66 | 1484.76 | 1482.75 | -2.01 | 19.8 |
| 18 | 104:112 | 1044.56 | 1044.56 | 0 | 14.4 |
| 19 | 119:126 | 779.33 | 779.34 | 0.01 | 17.7 |
| 20 | 135:140 | 651.35 | 651.35 | 0 | 9.7*** |
| 21 | 151:154 | 523.22 | 523.23 | 0.01 | 4.1 |
| 22 | 162:181 | 2191.97 | 2192.95 | 0.02 | 20.8 |
| 23 | 186:207 | 2330.18 | 2328.94 | -2.18 | 26 |
| 24 | 210:218 | 1000.5 | 1000.49 | -0.01 | 14.7 |
| 25 | 220:223 | 548.29 | 548.3 | 0.01 | 8.2 |
| 26 | 224:230 | 810.37 | 810.36 | -0.01 | 14.1 |
| 27 | 231:237 | 837.43 | ND | - | - |

Peptides 1-10 are the cysteine containing peptides predicted and, where indicated, identified after chymotrypsin digestion. Peptides 11-14 are the glutathionylated and native forms of peptides identified. Peptides 16-27 are the non-cysteine containing peptides predicted and, where indicated, identified after chymotrypsin digestion. The -sg indicates glutathionylated peptide. *These peptides containing cysteine 156 occur due to lack of cleavage at the 154:155 predicted chymotryptic cleavage site. ** These peptides containing Cys300 occur due to incomplete cleavage at the 305:306 predicted chymotryptic cleavage site. ND=Not Detected

**Table S2: RP/HPLC/MALDI-TOF MS Identification of peptides from trypsin/lysC digestion of $M^{pro}$ or monoglutathionylated $M^{pro}$ preparation**

| Peptide Number | Peptide From:To | $M_{r\ (calc)}$ | $M_{r\ (expt)}$ | Delta | R.T. |
|---|---|---|---|---|---|
| 1 | 13:40 (cys16,22,38) | 3446.56 | 3448.51 | 1.95 | 28.2 |
| 2a | 41:60 (cys44) | 2499.17 | 2500.14 | 0.97 | 24.6 |
| 2b | 41:61 (cys44) * | 2627.26 | 2628.23 | 0.97 | 23.8 |
| 3 | 77:88 (cys85) | 1452.71 | 1452.7 | -0.01 | 17.3 |
| 4a | 106:131 (cys117,128) | 3026.37 | ND | ND | ND |
| 4b | 106:137 (cys117,128) ** | 3726.76 | 3728.7 | 1.94 | 26 |
| 5 | 138:188 (cys145,156, 161 ) | 5561.38 | ND | ND | ND |
| 6 | 237:269 (cys265) | 3699.81 | 3701.75 | 1.94 | 29.2 |
| *7a* | 299:306 (cys300) | 993.41 | 993.41 | 0 | 15.1 |
| 7b | 299:306 (cys300)( pGlu) *** | 976.41 | 976.38 | -0.03 | 17.2 |
| 8 | 77:88 (cys-sg 85) | 1632.74 | 1632.71 | -0.03 | 13.5 |
| 8n | 77:88 (native cys 85) | 1327.66 | 1327.64 | -0.02 | 14.7 |
| 9 | 299:306 (cys-sg 300) | 1173.44 | 1173.42 | -0.02 | 10.9 |
| 9n | 299:306 (native cys 300) | 868.36 | 868.36 | 0 | 11.2 |
| 10 | 299:306 (pGlu) (cys-sg 300) | 1156.44 | 1156.4 | -0.04 | 12.7 |
| 10n | 299:306 (pGlu) (nat cys300)*** | 851.36 | 851.33 | -0.03 | 14 |
| 11 | 1:4 | 465.22 | 465.23 | 0.01 | 2.9 |
| 12 | 6:12 | 736.34 | 736.35 | -0.01 | 10.8 |
| 13 | 62:76 | 1695.87 | 1695.86 | -0.01 | 18.7 |
| 14 | 91:97 | 734.37 | ND | ND | ND |
| 15 | 132:137 | 718.39 | 718.37 | -0.02 | 8.8 |
| 16 | 189:217 | 3032.55 | 3032.54 | -0.01 | 25.6 |
| 17 | 218:222 | 734.38 | ND | ND | ND |
| 18 | 223:236 | 1613.8 | 1613.78 | -0.02 | 23.7 |
| 19 | 270:279 | 1130.54 | 1130.53 | 0.01 | 11.9 |

| | | | | | |
|---|---|---|---|---|---|
| 20 | 280:298 | 2121.09 | 2122.07 | -0.02 | 27 |

Peptides 1-7 are the cysteine containing peptides predicted and, where indicated, identified after trypsin/lysC digestion. Peptides 8-10 are the glutathionylated and native forms of peptides identified. Peptides 11-20 are the non-cysteine containing peptides predicted and, where indicated, identified after trypsin/lysC digestion. The -sg indicates glutathonylated peptide. *This peptide (41:61) was the result of incomplete cleavage at the 60/61 cleavage site. **This peptide was the result of incomplete digestion at the 131/132 cleavage site. ***These peptides are the result of the spontaneous deamidation that occurs with peptides containing an N-terminal glutamine, the retention times and molecular masses for this peptide were confirmed with the use of synthetic peptides that were run on RP-HPLC/MS. The retention times (RT) and molecular masses for the Cys300 peptides were confirmed with the use of synthetic peptides that were run on RP-HPLC/MALDI-TOF as native, alkylated or glutathionylated peptides. Peptide samples were analyzed without (-) and with (+) TCEP to remove glutathione moieties. Shown are the calculated native masses [$M_{r(calc)}$] and the experimental masses [$M_{r(expt0)}$]. The full TIC and 205 nm UV chromatograms for these analyses can be found in supplemental material (see Figure S5C-S5F in supplemental material). ND=Not Detected
